# DROP: Molecular voucher database for identification of *Drosophila* parasitoids

**DOI:** 10.1101/2021.02.09.430471

**Authors:** Chia-Hua Lue, Matthew L. Buffington, Sonja Scheffer, Matthew Lewis, Tyler A. Elliott, Amelia R. I. Lindsey, Amy Driskell, Anna Jandova, Masahito T. Kimura, Yves Carton, Robert R. Kula, Todd A. Schlenke, Mariana Mateos, Shubha Govind, Julien Varaldi, Emilio Guerrieri, Massimo Giorgini, Xingeng Wang, Kim Hoelmer, Kent M. Daane, Paul K. Abram, Nicholas A. Pardikes, Joel J. Brown, Melanie Thierry, Marylène Poirié, Paul Goldstein, Scott E. Miller, W. Daniel Tracey, Jeremy S. Davis, Francis M. Jiggins, Bregje Wertheim, Owen T. Lewis, Jeff Leips, Phillip P. A. Staniczenko, Jan Hrcek

## Abstract

Molecular identification is increasingly used to speed up biodiversity surveys and laboratory experiments. However, many groups of organisms cannot be reliably identified using standard databases such as GenBank or BOLD due to lack of sequenced voucher specimens identified by experts. Sometimes a large number of sequences are available, but with too many errors to allow identification. Here we address this problem for parasitoids of *Drosophila* by introducing a curated open-access molecular reference database, DROP (*<u>Dro</u>sophila* <u>p</u>arasitoids). Identifying *Drosophila* parasitoids is challenging and poses a major impediment to realize the full potential of this model system in studies ranging from molecular mechanisms to food webs, and in biological control of *Drosophila suzukii*. In DROP (http://doi.org/10.5281/zenodo.4519656), genetic data are linked to voucher specimens and, where possible, the voucher specimens are identified by taxonomists and vetted through direct comparison with primary type material. To initiate DROP, we curated 154 laboratory strains, 856 vouchers, 554 DNA sequences, 16 genomes, 14 transcriptomes, and 6 proteomes drawn from a total of 183 operational taxonomic units (OTUs): 114 described *Drosophila* parasitoid species and 69 provisional species. We found species richness of *Drosophila* parasitoids to be heavily underestimated and provide an updated taxonomic catalogue for the community. DROP offers accurate molecular identification and improves cross-referencing between individual studies that we hope will catalyze research on this diverse and fascinating model system. Our effort should also serve as an example for researchers facing similar molecular identification problems in other groups of organisms.

## Introduction

Building a knowledge base that encompasses ecology, evolution, genetics, and biological control is contingent on reliable taxonomic identifications. Molecular identification is commonly used in groups of organisms with cryptic species that are difficult to identify morphologically (Fagan-Jeffries et al., 2018; Miller et al., 2016; Novotny & Miller, 2014), for the molecular detection of species interactions (Baker et al., 2016; Condon et al., 2014; Gariepy et al., 2019; Hrček & Godfray, 2015; Hrcek et al., 2011), and for identification of species from environmental DNA samples (Shokralla et al., 2012). The accuracy of molecular identification, however, depends on the accuracy of identifications associated with sequences databased in existing online depositories (Fontes et al., 2021). The foundations of that accuracy are the voucher specimens which were sequenced and the collaboration of a taxonomic authority in the deposition of the sequence data.

GenBank serves as the most widely used sequence depository; however, deposition of sequences in GenBank, which is required by most peer-reviewed journals, does not require deposition of associated vouchers. The Barcode of Life Data System database (BOLD) (Ratnasingham & Hebert, 2007) explicitly aims to provide a framework for identifying specimens using single-locus DNA sequences (Hebert et al., 2003; Smith et al., 2005), and while these are associated with vouchers and metadata, the curation of these data is not consistently maintained by those submitting material. A recent study by Pentinsaari et al. (2020) showed misidentification in both databases caused by missteps in the protocols from query sequences to final determination.

Although the BOLD database function “BOLD-IDS” allows considerable database curation (e.g. flagging of misidentified/contaminated records), it also automatically includes sequences from GenBank, and may perpetuate the shortcomings previously mentioned since these cannot be curated from within BOLD. As such, the quality of sequences and the reliability of identifications obtained from BOLD-IDS can vary, and depends on the curation by taxonomists focusing on individual taxa (Meiklejohn et al., 2019). BOLD-IDS works well for taxa where qualified taxonomists have been involved with assuring data quality; some insect examples include beetles (Hendrich et al., 2015), butterflies (Escalante et al., 2010), geometrid moths (Hausmann et al., 2011, 2016; Miller et al., 2016), true bugs (Raupach et al., 2014), and microgastrine wasps (Smith et al., 2013).

Unfortunately, this is not the case of parasitoids (Insecta: Hymenoptera) of *Drosophila* flies (Insecta: Drosophilidae). There are vast numbers of *Drosophila* parasitoid sequences readily available in GenBank and BOLD, as these parasitoids and their hosts are important model organisms in biology. As of this writing, there are 88,666 nucleotide sequences deposited in GenBank for *Leptopilina heterotoma* (Thomson) and *L. boulardi* (Barbotin, Carton & Kelner-Pillault) (Hymenoptera: Figitidae) alone. However, less than 1 % of the identifications associated with these sequences have been confirmed by taxonomists or are associated with voucher specimens deposited in museum collections. With sequencing shifting from individual genes to genomes we risk that the identification problems will soon apply to whole genomes.

### Drosophila and their parasitoids

The phylogenetic and subgeneric structure within *Drosophila* and related genera is not yet fully resolved (O’Grady & DeSalle, 2018). Various subgenera, including *Scaptomyza*, *Zaprionus*, *Lordiphosa* and *Samoaia*, have been treated as both genera and subgenera, and researchers have yet to achieve consensus on these various hypotheses (O’Grady & DeSalle, 2018; Remsen & O’Grady, 2002; Yassin, 2013; Yassin & David, 2010). Species in *Drosophila* subgenera and genera closely related to *Drosophila* commonly share niche space and natural histories and, as a result, are often attacked by overlapping or identical groups of parasitoids. For instance, the invasive African fig fly, *Zaprionus indianus* Gupta is attacked by *Pachycrepoideus vindemiae* (Rondani, 1875) and *Leptopilina boulardi* (Pfeiffer et al., 2019; Santos et al., 2016), all of which have been recorded from *Drosophila*. Therefore, we also include these groups within the contents of DROP.

Parasitoids of *Drosophila* belong to four superfamilies of Hymenoptera (Chalcidoidea, Cynipoidea, Ichneumonoidea, Diaprioidea) which evolved parasitism of *Drosophila* flies independently. All the parasitoids known to attack *Drosophila* are solitary and attack either the larval or pupal stage; in both cases, they emerge from the fly’s puparium (Carton et al., 1986; Prévost, 2009). The known *Drosophila* larval parasitoids belong to two families (Table 1), Braconidae (including the genera *Asobara, Aphaereta, Phaenocarpa, Tanycarpa, Aspilota, Opius*) and Figitidae (*Leptopilina, Ganaspis, Leptolamina, Kleidotoma*); all are koinobionts that allow the host to continue development while the parasitoid grows within it. The known *Drosophila* pupal parasitoids belong to three other families (Table 1), Diapriidae (*Trichopria, Spilomicrus*), Pteromalidae (*Pachycrepoideus, Spalangia, Trichomalopsis, Toxomorpha*) and Encytidae (*Tachinaephagus*); they are all idiobionts that terminate host development immediately. Host-specificity across the *Drosophila* parasitoids is poorly characterized—while some can parasitize other families of Diptera (e.g., *Aphaereta aotea*) (Hughes & Woolcock, 1976), most are thought to be limited to *Drosophila* hosts.

**Table 1:**
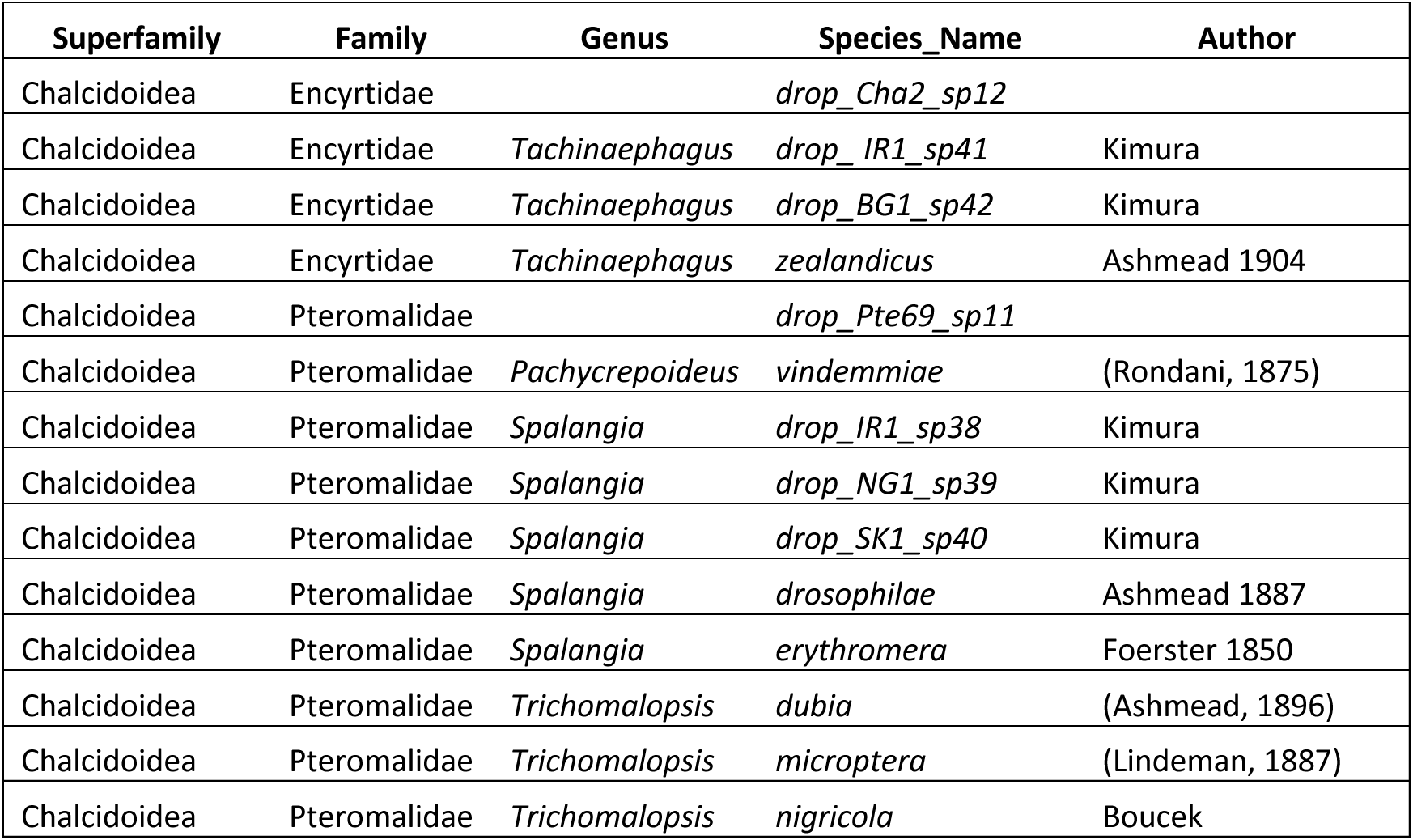
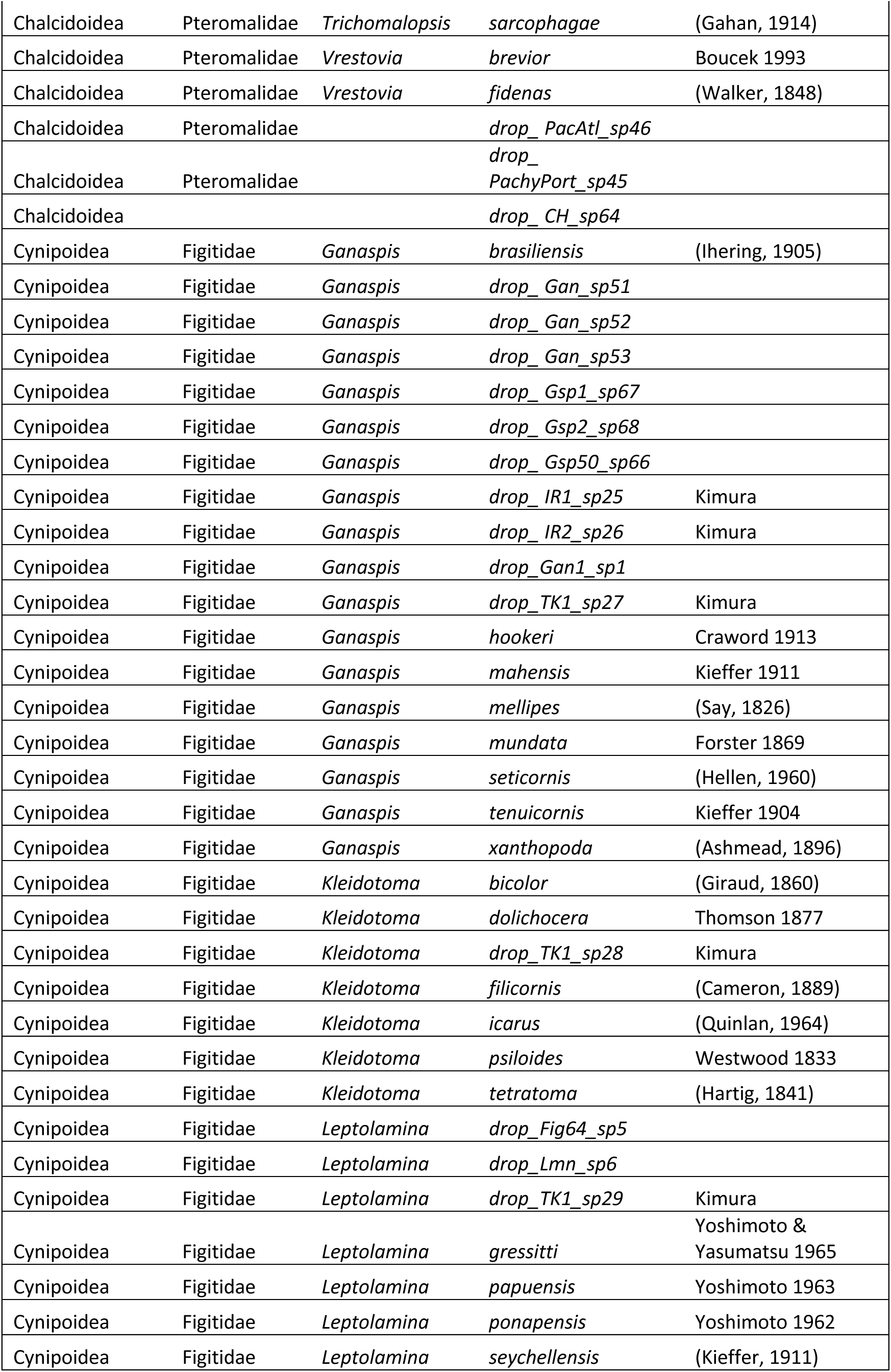
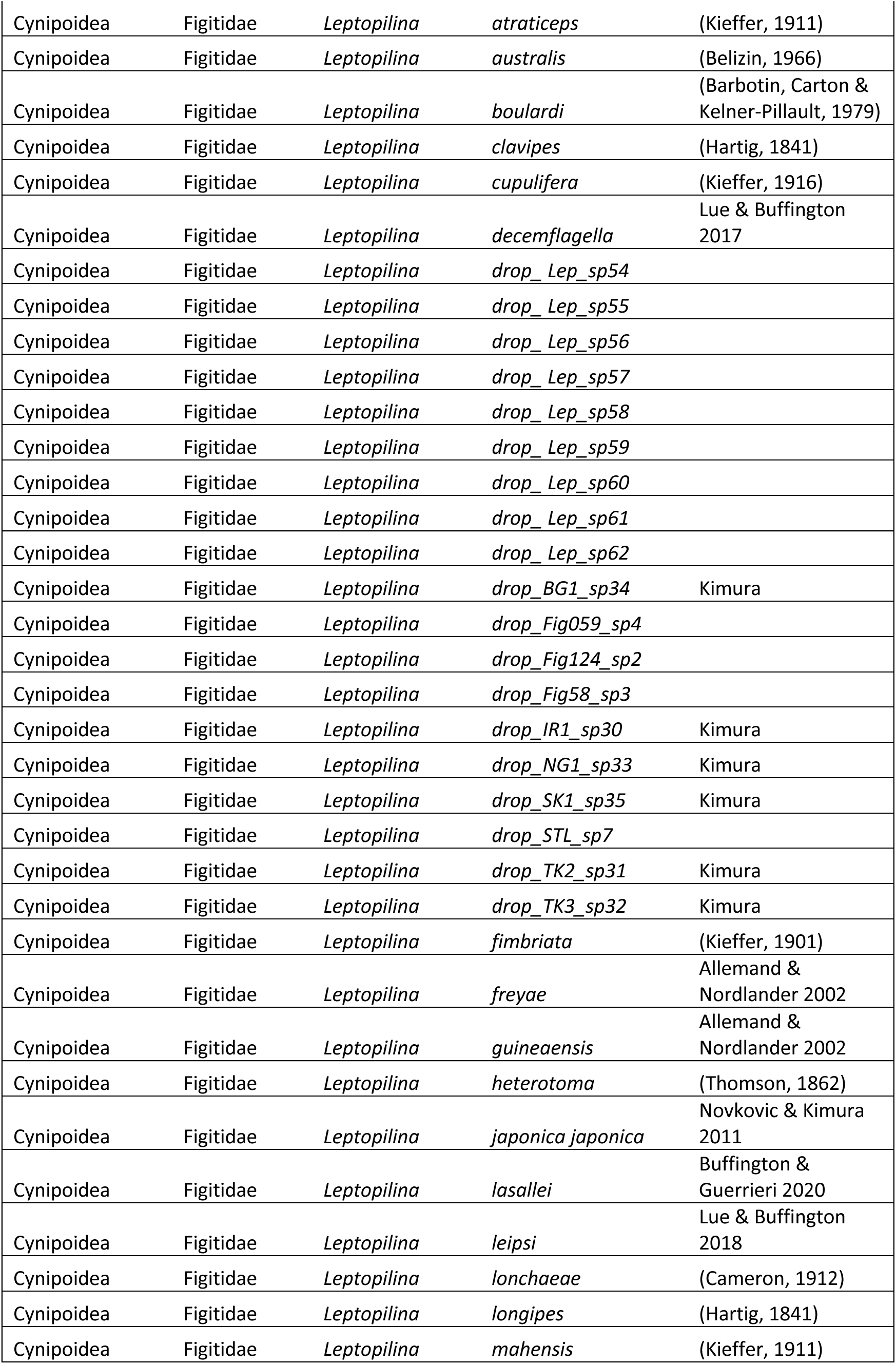
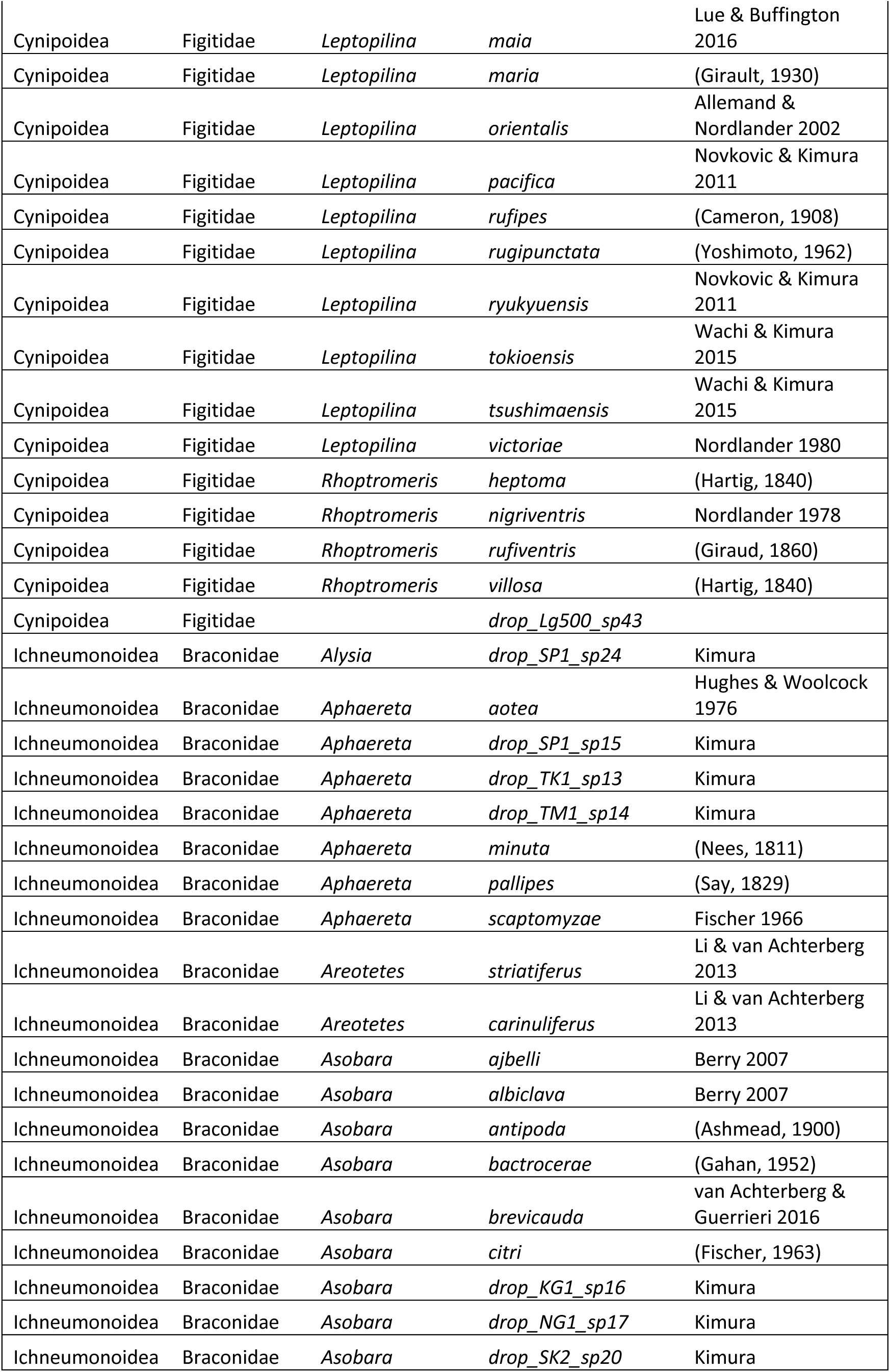
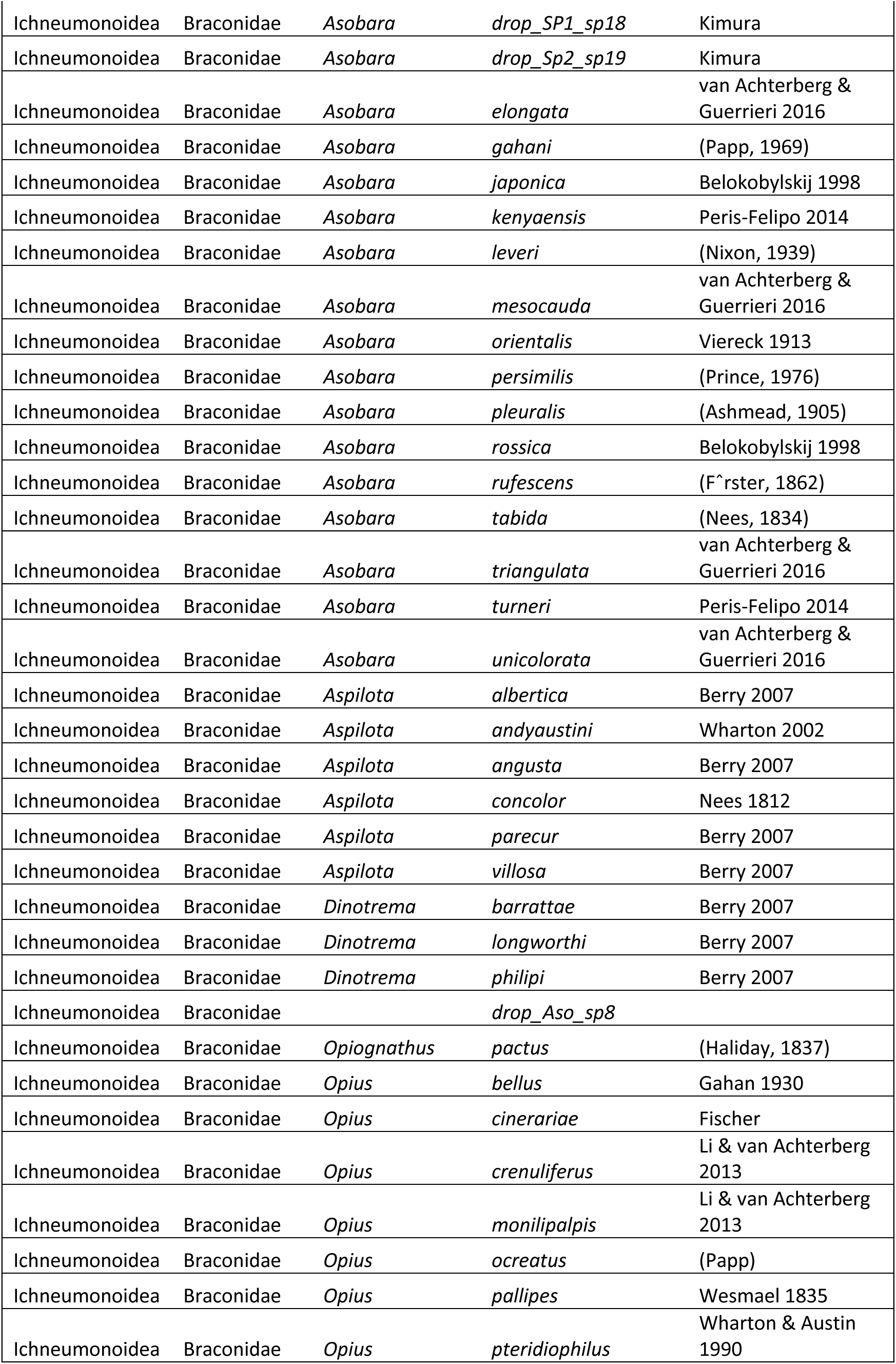
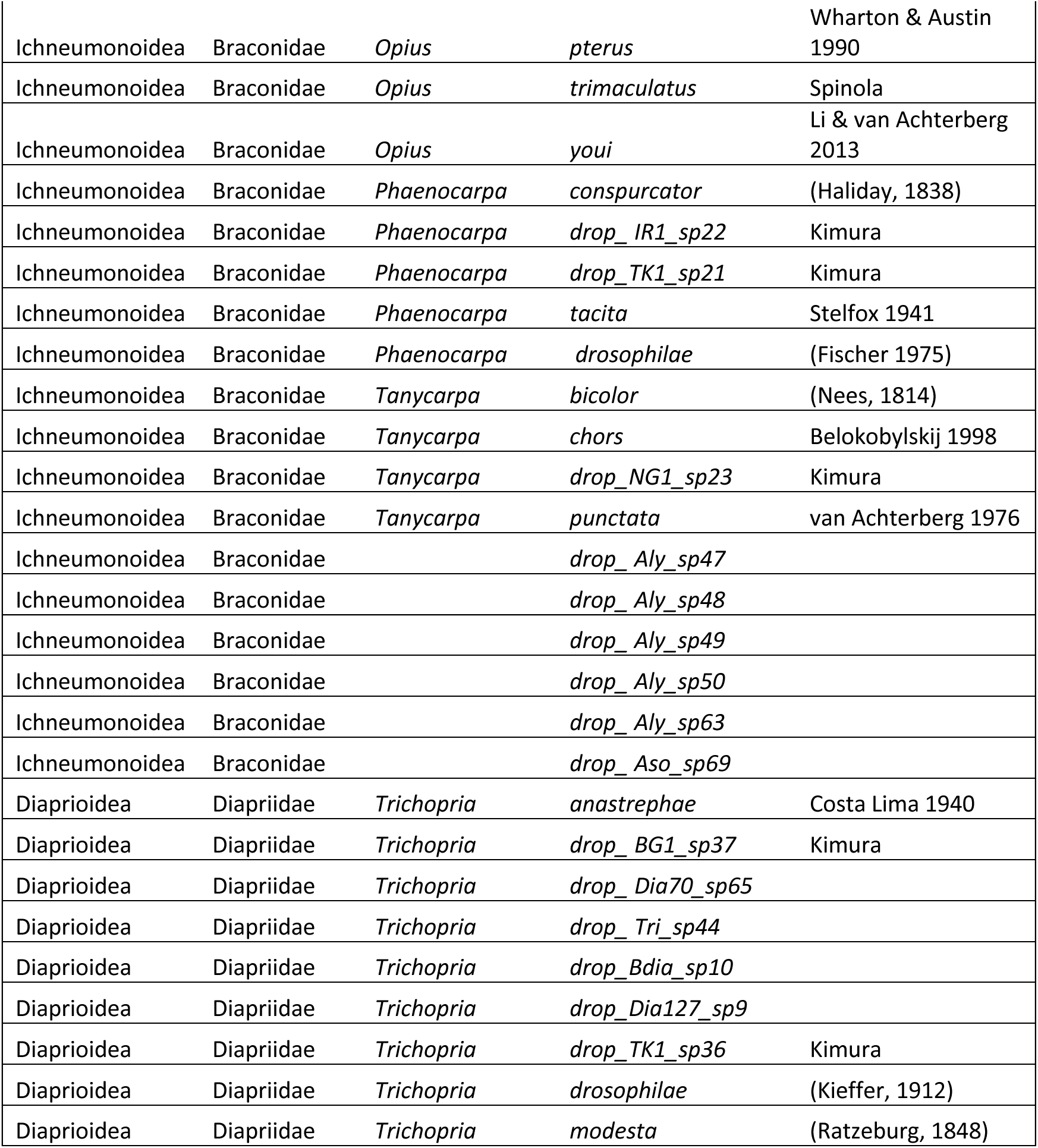
List of species and provisional species included in DROP. For additional taxonomic details, see DROP.

There are around 4000 described species of Drosophilidae, and *Drosophila* contains more than a third of the family’s described species (O’Grady & DeSalle, 2018). By contrast, although parasitic wasps are generally a species-rich group (Dolphin & Quicke, 2001; Quicke, 2015), the most recent catalogue of parasitoid species that attack *Drosophila* lists only 50 described species (Carton et al., 1986). This disparity suggests that the diversity of parasitic wasps attacking *Drosophila* is severely underestimated, an assertion supported by the results presented here. This is largely a consequence of the challenging nature of parasitoid taxonomy, in which morphological identification is intractable for many species, and the fact that taxonomic specialists are greatly outnumbered by the species they study.

Currently, only a few biological study systems have been characterized in sufficient breadth and depth to allow researchers to connect various levels of biological organization, from molecular mechanisms to food webs of interacting species. Parasitoids of *Drosophila* represent one such system (Prévost, 2009). Moreover, the practical feasibility of rearing parasitoids of *Drosophila* under laboratory conditions has led to a number of fundamental discoveries in ecology (Carton et al., 1991; Terry et al., 2021), evolution (Kraaijeveld & Godfray, 1997), immunology (Kim-Jo et al., 2019; Nappi & Carton, 2001; Schlenke et al., 2007), physiology (Melk & Govind, 1999), symbiosis (Xie et al., 2011, 2015), behavioral science (Lefèvre et al., 2012) and other fields. In contrast to this large body of laboratory studies, basic natural history of *Drosophila* parasitoids, especially their species richness is little known (Kimura & Mitsui, 2020; Lue et al., 2018). Addressing this knowledge gap is especially pressing given current efforts to use parasitoids in biological control efforts, such as those of the invasive pest spotted wing *Drosophila*, *Drosophila suzukii* (Abram et al., 2020; Daane et al., 2016; Giorgini et al., 2019; Wang et al., 2020 a&b).

Properly executed molecular identification has the potential to be much more efficient for the majority of researchers, and many laboratory strains are commonly identified using DNA sequences alone. While it is practical for researchers to assign species names based on a match to sequence records in genetic databases, this practice often causes a cascade of inaccuracies. To illustrate the extent of the problem, we present the example of *Ganaspis,* a genus of parasitoids commonly used in laboratories that includes both superficially indistinguishable species with highly divergent sequences that are often treated as conspecific, as well as specimens with identical sequences identified under different names (Figure 1).

**Figure 1:**
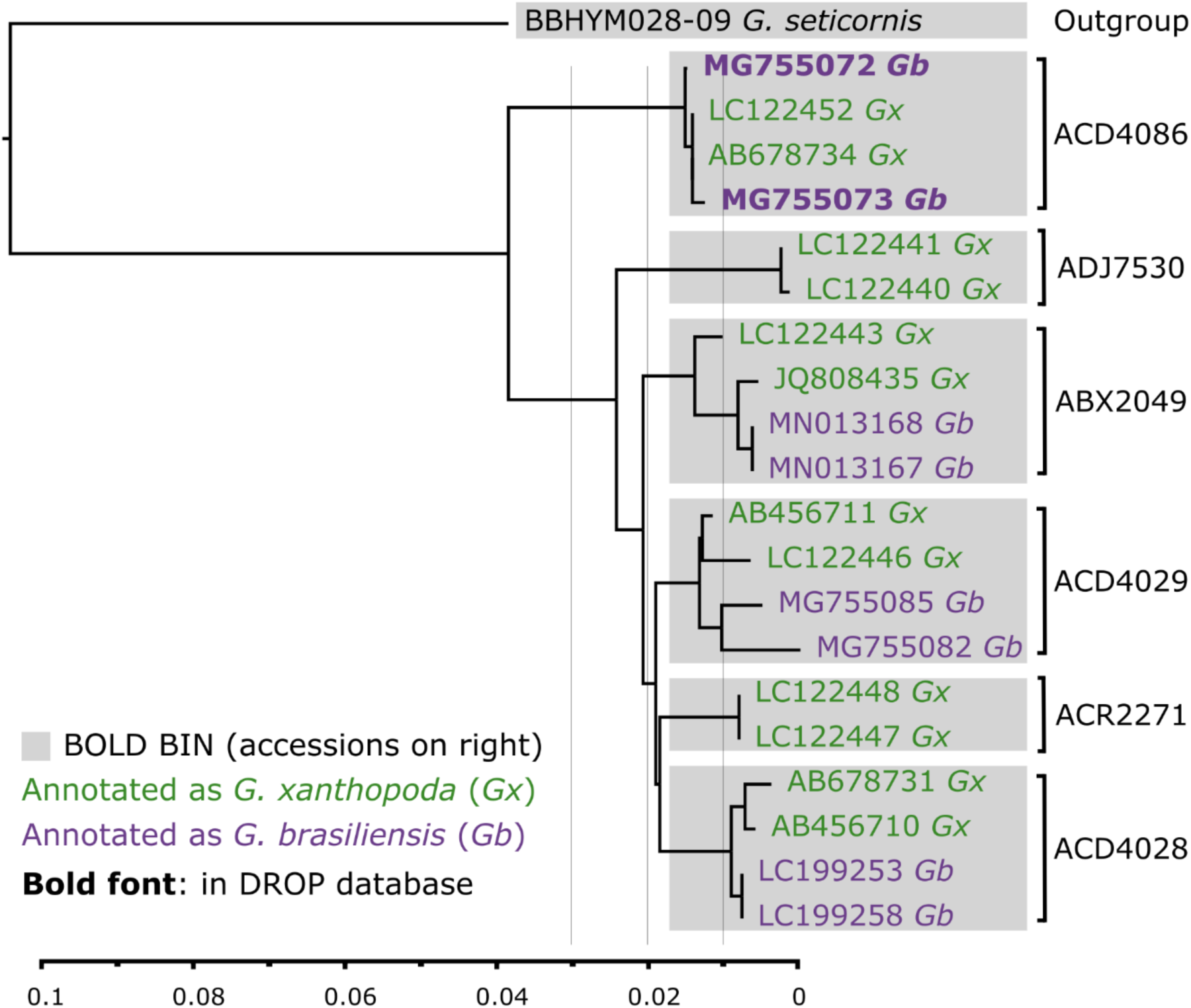
An example of difficulties of molecular identification demonstrated on *Ganaspis xanthopoda* and *G. brasiliensis*. Only two sequences (in bold text) can be reliably used for identification and are included in DROP database. To select the sequences, we searched the BINs associated with the organism’s name “*Ganaspis xanthopoda”* (green) or *“Ganaspis brasiliensis”* (purple) in BOLD. From each BIN, two sequences from each species were selected to build a neighbor-joining tree (bottom axis indicated % genetic divergence). There was a total of 6 BINs (gray boxes) in this sequence complex. Of these, 4 BINs contained both species names and without examination of vouchers, identification would be impossible. In DROP, vouchers from two sequences, **MG755073** and **MG755072**, were deposited in CNR-IPSP (Table S2), examined by taxonomists and identified as *G. brasiliensis*. These two COI sequences can now be used to reliably identify *G. brasiliensis*. For *G. xanthopoda,* there were no available vouchers or reliable sequences that passed DROP standards to use for identification. Species delimitation between *G. brasiliensis* and *G. xanthopoda* is convoluted, varies according to arbitrary % genetic divergence (gray vertical lines), and needs an integrative taxonomic revision.

### Aims

To address these issues, we introduce a newly curated molecular reference database for *<u>Dro</u>sophila* <u>p</u>arasitoids —DROP— in which sequences are either linked to voucher specimens identified by taxonomists or have a traceable provenance (Figure 2). The first aim of DROP is to provide a reliable DNA sequence library for molecular identification of *Drosophila* parasitoids that enables cross-referencing of original taxonomic concepts with those of subsequent studies. We pay special attention to live parasitoid strains which are available for future experiments. The second aim is to standardize and expedite the linkage between specimens and available sequence data; we place a premium on museum vouchers as they allow for repeatable scientific research. In DROP, this goal is facilitated through a consolidated digital infrastructure of data associated with laboratory strains, offering the opportunity for researchers to re-examine past experimental results in a permanent context. The third aim is to provide an up-to-date catalogue of the diversity of *Drosophila* parasitoids as a foundation for advancing the understanding of their taxonomy. Finally, the fourth aim of DROP is for our collaborative effort to serve as an inspiration to communities of researchers studying other groups of organisms who are experiencing difficulties with the reliability of molecular reference databases.

**Figure 2:**
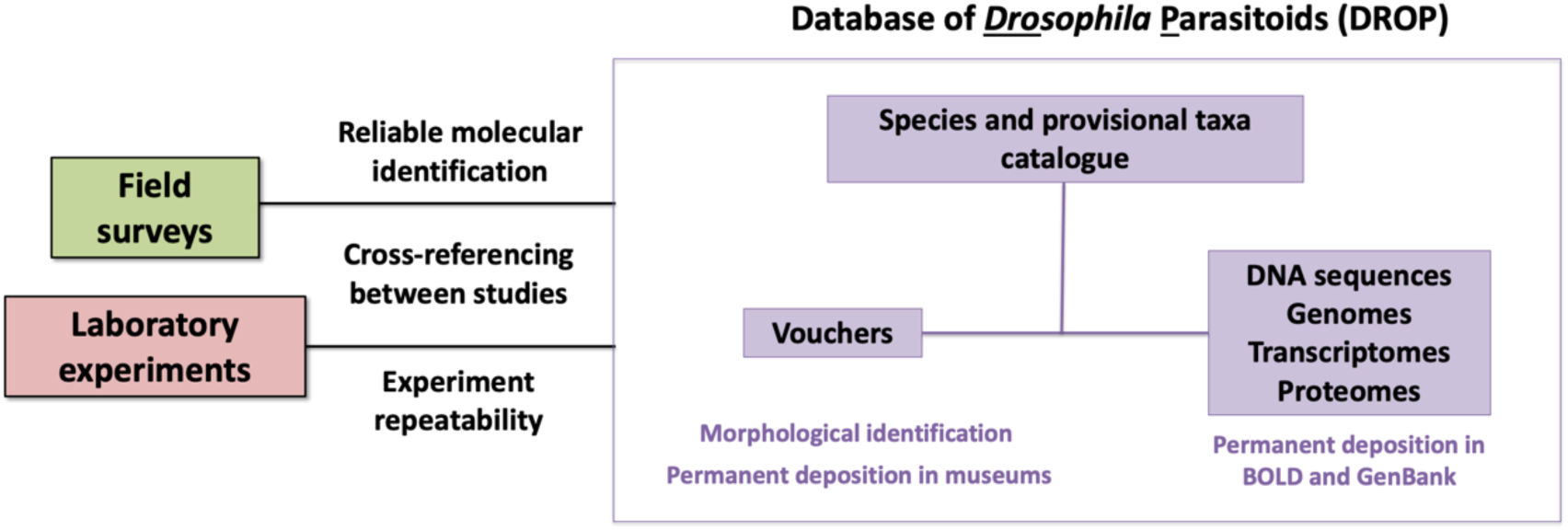
Concept of a centralized, vetted, curated database for *<u>Dro</u>sophila* <u>P</u>arasitoids (DROP) we developed. First, we provide a species and provisional species catalog with correct taxonomy. Second, to provide a reliable genetic reference library, we link genetic data (DNA sequences, genomes, transcriptomes, proteomes) to a voucher connected to the species catalog. Third, we link the two primary sources of data (field surveys and laboratory experiments) by requiring a permanent deposition of vouchers and sequences in order to be included in DROP.

## Materials and Methods

### Data sources

To assemble the DROP database, we targeted 20 wasp genera that potentially parasitize frugivorous *Drosophila* species. We compiled DNA sequence and voucher data from four sources: 1) museum collections, 2) publications, for which we selected the reference with taxonomist or parasitoid biologists as coauthors to ensure reliable species identity, 3) molecular biodiversity inventories publicly available in BOLD and GenBank, for which we managed to secure inspection of the vouchers by taxonomists, and 4) a sequencing and taxonomic inventory of laboratory strains we conducted.

We first gathered species information into a catalogue of *Drosophila* parasitoid species (Table 1) from 216 references (see DROP database reference table) and 36 institutes (Table S2). To ensure reliable names for nominal species (sequences identified by a species name) in our database, we confirmed their taxonomic validity using the Ichneumonoidea 2015 digital catalogue (Yu et al., 2016) and Hymenoptera Online (HOL; http://hol.osu.edu/), both of which are curated by taxonomic experts. To obtain reliable molecular identification data, we harvested 8,298 DNA sequences from GenBank and BOLD (all compiled in BOLD as DS-DROPAR dataset <u>dx.doi.org/10.5883/DS-DROPAR</u>). As of writing, these sequences represented 445 Barcode Index Numbers (BINs – a form of dynamic provisionary taxa in BOLD, more detail in Ratnasingham & Hebert 2013), and 211 named taxa.

The majority of the harvested sequences were Braconidae (6690), Diapriidae (967), Figitidae (622), and Pteromalidae (19). Because of the concerns with generic databases (noted above and in Figure 1 and Table S1), we assembled a list of sequences with valid species names that could either be traced back to vouchers examined by taxonomists or were referred to directly in publications authored by a recognized expert in the relevant taxon group. We then cross-checked species names with their corresponding BINs in BOLD and flagged potential conflicts between species names and BINs (Table S1).

A core goal of DROP besides that of a tool for biodiversity research is to function as a platform that accommodates *Drosophila* parasitoids kept in laboratory strains (for experimental work) or cultures in quarantine facilities (for biological control applications). So far, there has been a lack of a coherent and reliable means of verifying identification of species kept in laboratory settings, which can be a serious problem. Since lab cultures are routinely contaminated by neighboring cultures (e.g., through escapees), one species may be displaced by another even under a vigilant eye.

For lab and quarantine lines in DROP, we deposited DNA extractions and vouchers in the National Insect Collection, National Museum of Natural History, Smithsonian Institution (USNM; Washington, DC, USA). During their initial assembly of DROP, laboratory OTUs (operational taxonomic unit) were designated by their strain name; most laboratory strains can be associated with provisional species, but some cannot yet be assigned. Three females and three males of each strain were dry-mounted and individually assigned a USNMENT ‘QR code’ specimen label as representative vouchers. For each molecular voucher, three legs from a female wasp were removed for DNA extraction and sequencing (Supplementary Methods for details), and the rest of the body was assigned a USNMENT specimen label and preserved for morphological identification. Both DNA extraction and vouchers were entered into the database and uploaded to BOLD (DROP project: DS-LABS <u>dx.doi.org/10.5883/DS-LABS</u>) with an associated GenBank ID.

Where possible, we identified OTU strains using a combination of morphological and sequence data, and characterized provisional species or species clusters using neighbor-joining trees (Figure S1) based on the COI gene sequences (Supplemental material). For establishing BIN limits in the context of DROP, we have adopted an initial percent cutoff at 2%. We acknowledge that 2% genetic diverge cutoffs (or BINs) are unlikely to work well across range of widely distributed species (Lin et al., 2015). But as Ratnasingham & Hebert (2013) pointed out, 2% is a good starting point for many taxa, also it may need to be adjusted as more samples are acquired and compared. Note that we use the term “OTU” as a general and neutral designation encompassing described species, provisional species, undescribed species, and cryptic species.

### Drosophila parasitoid database—DROP

To compile the above information, we built a simple Structured Query Language (SQL) database in sqlite3 format using SQLiteStudio (step by step user instruction in supplemental material). Sqlite3 is a cross-platform format which can be also be opened using a number of other programs. There are eight linked tables in the database — species, strain, voucher, sequence, genome, transcriptome, proteome and reference — along with additional tables for linking these to reference table (Figure S2). The database incorporates all sample fields used by BOLD for compatibility and includes a number of new fields to accommodate a catalogue of *Drosophila* parasitoid species, laboratory strain information, and links from the DROP database to BOLD and GenBank records.

DROP is available on Zenodo (http://doi.org/10.5281/zenodo.4519656) for permanent deposition and version control. In addition to the main database, the Zenodo repository includes additional files to facilitate easy use of the database. These files include: 1) the reference database in comma-separated text (.csv) and FASTA format ready to be used for molecular identification; 2) a species catalogue with taxonomic information; and 3) a list of laboratory strains with confirmed molecular vouchers. DROP will be continued to be curated and maintained by C-HL at the Zenodo repository and sequences generated in the future will also be deposited in BOLD (DROP project). If the curator changes, this will be announced in the README.md file in Zenodo repository. As the database relies on vouchers, we will aim for it to be continued to be maintained by taxonomists with direct access to museums.

### Species, provisional species, and OTU designations

In addition to the inherent value of a formal taxonomic name, a reliable provisional taxon label can also be used for exchanging scientific information and conveying experimental results among researchers (Schindel & Miller, 2010). Based on the amount of sequence divergence between described species, we observed what appears to be a significant number of provisional OTUs in the initial dataset we compiled. Furthermore, among the data linked to a valid species name, some of these provisional OTUs are actively being used in research and have sequences available to the public. We therefore provide a list of provisional species (potential new species) with their molecular vouchers.

We use the following designation format for OTUs that refer to a provisional species: “Drop_strainX_sp.1” or, when no other information is known, “DROP_sp.1”. Where possible, these OTUs are linked to a voucher USNM specimen label number. If the genus of the OTU is known, the “Drop_Leptopilina_sp.1” format is followed. These designations can facilitate species identification as well as discovery and description of new species without compromising the existing taxonomy of the described OTUs in question. As more complete species descriptions become available, this provisional species framework can be updated while keeping the link to previous provisional species name through deposited vouchers.

## Results

### Overview of DROP

We catalogued 183 OTUs in the DROP database with 114 described species of *Drosophila* parasitoids and 69 provisional species (Table 1). In total, we documented 154 laboratory strains (Table S3), and 853 vouchers from 36 institutions (Table S2). Among the described species, 98 have voucher information, of which 61 are traceable to type specimens, including 45 to holotypes (i.e., specimen used to root a name to the taxonomic author’s concept of the species). *Leptopilina* is represented by the highest number of species with 45 OTUs, followed by *Asobara* with 26 OTUs. Within the 154 catalogued lab strains, 86 were actively being used in ongoing research (i.e., a live strain being cultivated). These strains represent 39 OTUs: 11 described species and 28 provisional species (Table S3, Figure S1).

### Molecular Vouchers

So far, DROP includes 545 DNA sequences and links to 16 genomes (Table S4), 14 transcriptomes (Table S5), and 6 proteomes (Table S6). From the total of 8298 DNA sequences (BOLD dataset: DS-DROPAR) collected from public databases, only 322 sequences (less than 4% of available sequences) satisfied the reliability criteria we imposed for molecular vouchers to be included in DROP (see Materials and Methods). The DS-DROPAR dataset <u>dx.doi.org/10.5883/DS-DROPAR</u> initially referred to 211 taxon names, but only 52 names were valid, linked to vouchers, or linked to a publication with evidence that the specimens had been identified by taxonomists. The remaining 223 of 545 DROP DNA sequences were generated by DROP project (datasets: DS-LABS <u>dx.doi.org/10.5883/DS-LABS</u> and DS-AUSPTOID <u>dx.doi.org/10.5883/DS-AUSPTOID</u>) and came from 121 OTUs (101 lab strains and 12 provisional species).

The DROP database is largely made up of standard barcode COI sequences (349 sequences), which includes 77 OTUs: 43 described species and 33 provisional species. We aimed to supplement COI with secondary markers (28SD2, 18S, ITS2) when possible, resulting in an additional 120 sequences from 26 OTUs: 15 described species and 11 provisional species. There are currently 19 OTUs that have sequences from more than one genetic marker.

### Species Delimitation in Laboratory Strains

We used 298 COI sequences to resolve the identification of each laboratory strain, and where possible, indicated potential species clusters (Fig. S1 and Table S3). Using a fixed 2% divergence cutoff, a total of 31 lab strain OTUs were assignable to a valid species name, and the remaining 70 strain OTUs were assigned to a provisional species. The taxonomic status of several of these provisional species is also being investigated using an integrative taxonomic approach involving morphological identification, genomic data, or other genetic data.

## Discussion

In this paper, we introduce and describe a free and open-access database for the reliable molecular identification of *Drosophila* parasitoids. The guiding principle of DROP is data credibility, based on the prerequisite that genetic data are explicitly associated with voucher specimens and taxonomic concepts of the original authors (Troudet et al., 2018). When incorporating information from public genetic databases, we included only sequences that have passed our filtering protocol. This protocol ensures each entry is associated with a valid scientific name, provisional name, or consistently applied OTU designation that can be used to integrate genetic and organismal data from independent studies.

The following discussion expands on the utility of DROP and how we hope it will benefit molecular species identification, connect research from various disciplines, support biological control applications, and serve as a long-term molecular voucher repository and clearinghouse for vetted data. We also provide specific guidance for users how best to refer to DROP in their publications to allow cross-linking between studies.

### Molecular (mis-)identification

We observe that 17% of the described *Drosophila* parasitoid OTUs in BOLD and GenBank (dataset: DS-DROPAR) are associated with more than one BIN; these are examples of BIN-ID conflict. Roughly half of these OTUs are used as lab strains. This latter observation is disturbing, because it demonstrates that the criteria used to differentiate and reference species in active research programs are clouded. For example, BIN-ID conflicts were observed in the *Drosophila* parasitoids *Ganaspis brasiliensis* (Ihering) and *Asobara japonica* Belokobylskij (Table S1), both of which are in active use in numerous research programs (e.g. Moreau et al., 2009; Nomano et al., 2017; Reumer et al., 2012; Wang et al., 2020a & 2021) as well as in biological control efforts against the invasive *D. suzukii* (e.g. Abram et al., 2020; Daane et al., 2016; Giorgini et al., 2019). All the BINs from *G. brasiliensis* carry the name *G. xanthopoda* (Figure 1). In such instances, assigning an identification by matching specimens to barcode records in the genetic database is problematic, as two names are applied to the same BIN. If sequences comprising the BIN are not linked to a voucher that can be examined, teasing apart the two names and how they are applied is impossible. Applying explicit, consistent criteria for species determination ensures that experimental results can be reliably repeated, and that any potentially novel observations will not be explained away as artifacts of identification. DROP addresses these concerns by linking reliable reference sequences and vouchers for *G. brasiliensis* (Figure 1) between different studies: one with reference to the morphological description (Buffington & Forshage, 2016) and the other with reference to the genome (using voucher specimens from the morphological study; Blaimer et al., 2020).

We were not able to resolve all conflicts between BIN and species identity, for one or more of the following three reasons: First, many records lack reliably identified vouchers and have often been themselves used for molecular identification, proliferating errors. Second, in some cases, it is not possible to verify whether the genetic differences among BINs represent different species or simply intraspecific genetic variation (Bergsten et al., 2012), because BINs themselves are not a species concept. The only solution to this problem is to derive original sequence data from type specimens (which is often either impractical or impossible for a number of technical reasons), or from specimens whose conspecificity with the types has been corroborated. Since species boundaries are always subject to testing, additional specimens from multiple collecting events (ideally representing different seasons and geographic regions) may help provide the additional data to circumscribe a given species’ limits. The third difficulty in resolving BIN-ID conflict derives from the data themselves: Although the mitochondrial COI gene is the locus most frequently chosen for identification of insects and other animals, its effectiveness varies among insect groups (Brower & DeSalle, 2002; Gompert et al., 2008; Lin & Danforth, 2004). In part, this derives from gene-tree/species-tree conflict as a function of mitochondrial DNA introgression (Gompert et al., 2008; Klopfstein et al., 2016), parthenogenesis (Reumer et al., 2012), and/or *Wolbachia* infection (Ferrer-Suay et al., 2018; Wachi et al., 2015; Xiao et al., 2012), any of which may lead to complications in species delimitation using mitochondrial loci. Ideally, studies should apply multiple loci, genomes, and comparative taxonomic data to clarify species boundaries. As *Drosophila* parasitoids are often maintained in laboratory cultures, it is also possible to use mating experiments to explore species boundaries under the paradigm of the biological species concept (Seehausen et al., 2020).

### DROP as a taxonomic tool

DROP offers an empirical platform for species discovery and a useful tool for taxonomic research. The fact that the number of BINs reported here exceeds the number of described species (Table S1, Figure S3) highlights the need for taxonomic work. But such work cannot proceed on the basis of BINs or barcodes, but requires integrative taxonomic approach employing a combination of molecular and morphological data. Describing new species on the sole basis of a barcode or BIN, without the benefit of independent character data, should, in general, be avoided (Meier et al., 2021). It risks creating nomenclatural synonymy if it is later determined that a sequence can be attributed to a specimen that bears a valid, available name. Moreover, BINs are based on distance analyses which, by definition, are incompatible with diagnoses per se (Ferguson, 2002; Prendini et al., 2002; Goldstein & DeSalle, 2011). Therefore, in taxonomic treatments, it is critical to clarify the range of applicability of a given BIN and its overlap with a taxonomic name (see example in Figure 1). DROP allows cross-linking between studies and therefore provides researchers with valuable tools for taxonomic revisions, including the means of discovery, corroboration, and description of new species.

### How to use DROP to ensure cross-linking between studies and reliable molecular identification?

Public genetic databases have adopted a longstanding convention in treating undetermined OTUs and sequences, referring to provisional species with numbers, as for example “sp. 1*”*, and these are rarely linked to vouchers. For OTUs designated as provisional species, DROP enables cross-indexing of specimens, sequences and references between any studies (ecological, taxonomic, evolutionary, genetic, etc). The best way to ensure cross-linking is depositing a voucher in DROP, together with a sequence or genome from the same individual (or individual from the same strain or series). For example, one can write:

*Provisional species “drop_Gan1_sp.1” refers to voucher USNMENT01557320 deposited in the USNM, Washington DC, COI sequence (DROP sequence_id: 2, BOLD Process ID: DROP143-21), 28SD1 sequence (DROP sequence_id: 289), and 28SD2 sequence (DROP sequence_id: 303)*.

Similarly, laboratory strains can be reported in the same way, just adding the DROP lab strain_id. It is important to periodically recheck identification of laboratory strains as cultures are easily cross-contaminated, and deposit vouchers of laboratory strains associated with experiments to DROP. In the future, when e.g. “drop_Gan1_sp.1” is described as a new species with a formal specific epithet, DROP curator will update the species status and holotype information while keeping this provisional species name as an informal “synonym.”

A weaker and thus much less preferred way of cross-linking is to state in the study that the identification of organisms was performed based on molecular identification match of a sequence to DROP sequences. This is the only available option for environmental DNA studies. For example, one can write:

*Provisional species “drop_Gan1_sp.1” was identified based on 99.9% blast match of COI to DROP sequence_id: 2 (BOLD Process ID: DROP143-21)*.

DROP deposition in Zenodo allows referencing of DROP either through general doi (the doi we use throughout this paper), which takes the user always to the latest database version, or through a doi specific to DROP version. When referencing DROP please primarily cite this paper, but for reproducibility it is also good practice to include doi of the specific DROP version used.

There are two basic ways of molecular identification which should ideally be used in combination: sequence matching (blast), and tree-building methods which investigate membership to a cluster. Further, there are a number of decisions to be made with each method, concerning locus (or loci) and thresholds. DROP leaves these decisions up to the users, only provides raw sequences or links to them. Practically, the choice of loci is currently mostly limited to COI, but in the future it is likely that molecular identifications will be based on multiple loci or whole genomes. Over time we will also get a better idea about what thresholds are more appropriate than a fixed 2% cut off. For rarer parasitoid genera which attack also other hosts besides *Drosophila* (e.g. *Opius*, or *Spalangia* wasps) we suggest caution in the identification using only DROP sequences as DROP does not include all sequences from these genera, but just from species which are already known to attack *Drosophila*.

### From molecular mechanisms to ecosystem structure

The use of molecular tools in insect biodiversity studies has gradually expanded from barcoding single individuals to metabarcoding large environmental samples representing entire food webs (Jeffs et al., 2020; Littlefair et al., 2016). *Drosophila* and their parasitoids are among the few systems that currently allow us to explore thoroughly the mechanisms of species interactions at scales ranging from the molecular to the ecological. Here, we highlight two examples where information compiled in DROP enables the study of the *Drosophila*-parasitoid system across multiple levels of biological organization:

DROP includes a DNA reference library of Australian *Drosophila* parasitoids (dataset: DS-AUSPTOID <u>dx.doi.org/10.5883/DS-AUSPTOID</u>) that connects laboratory experiments and field research. Molecular vouchers of both hosts and parasitoids were collected along altitudinal gradients in the rainforest of northern Queensland, Australia (Jeffs et al., 2021). With this DNA reference library, researchers can detect interactions between *Drosophila* and their parasitoids using PCR-based approaches and parasitized pupae (Hrcek & Godfray, 2015; Jeffs et al., 2020). Surveying host-parasitoid interactions in this way will improve our understanding of how environmental change alters the structure of host-parasitoid networks (Morris et al., 2014; Staniczenko et al., 2017; Tylianakis et al., 2007) by accelerating data collection in the field. In addition, JH established lab cultures of both hosts and their parasitoids from the same Australian sampling sites with the aim of conducting laboratory experiments (e.g. Thierry et al., 2021). Molecular vouchers of the lab strains were then submitted to DROP as a reference database (datasets: DS-LABS <u>dx.doi.org/10.5883/DS-LABS</u>) to ensure that criteria for species determination were applied consistently—and will continue to be applied consistently— between the natural community studies and the laboratory experiments.

The presence of a foundational DNA reference library and species catalogue in DROP will enable the process of exploring parasitoid biodiversity to become more efficient. For example, DROP includes molecular vouchers from *Drosophila* parasitoids that were collected across seasons and along latitudinal gradients in the eastern United States (Lue et al., 2016, 2018). These data proved to be extremely useful for identifying species in a more recent exploration of native parasitoid biodiversity across North America (e.g., Abram et al., 2020). There are additional uses for DROP: curated specimen collections may be used to document species distributions, phenology, understand micro-evolutionary patterns, observe the effects of climate change, and detect and track biological invasions (Funk, 2018; Schilthuizen et al., 2015; Tarli et al., 2018).

### Taxonomic accuracy for biocontrol studies

Unfortunately, the history of biological control includes many examples of misidentifications that have resulted in failures to employ or establish the expected control agent, thus hindering eventual success (Buffington et al., 2018; Rosen, 1986; Huffaker et al. 1962). In the context of biological control research on *Drosophila* pest species, a simple, reliable, and rapid identification tool for their natural enemies is essential (Wang et al. 2020b). By anchoring the criteria for determining identities of organisms being considered for biological control programs, DROP annotation enables the direct examination of centers of origin for parasitoid species, their co-occurrence with natural enemies, and the optimal timing for potential introductions of such enemies (Abram et al., 2020; Daane et al., 2016; Girod et al., 2018a and b; Kimura, 2015; Mitsui et al., 2007). Because most sequences from DROP are already vetted for reliability, they can be used to identify biological control agents rapidly, before or after being brought into quarantine facilities for safety and efficacy testing. This will decrease the risk of non-target ecological impacts arising from misidentifications and facilitate regulatory review for releases of effective and specific natural enemies.

In addition to species identification, reference sequences from DROP may be used as a starting point to create species-specific primers for the accurate identification of parasitoids, design multiplex PCR assays that rapidly distinguish species in natural or agricultural ecosystems (Ye et al., 2017), and apply high-throughput molecular identification diagnostics (Fagan-Jeffries et al., 2018).

### Long-term molecular voucher preservation

During the curation of DROP, we found that holotype specimens were missing from museums for several iconic *Drosophila* parasitoid species: *Asobara tabida* (Nees von Esenbeck), *Leptopilina clavipes* (Hartig), and *Leptopilina longipes* (Hartig). This is not uncommon and impedes future taxonomic revisions regardless of whether or not molecular data are used. To avoid contributing to this problem, DROP uses museums as depositories for ensuring that sequenced vouchers of both described species and provisional species are permanently stored. In order to stabilize nomenclature, we further advocate the designation of neotypes (a replacement specimen for a missing holotype or type series) that have museum-vouchered DNA barcodes and additional genomic extractions in storage.

Natural history museums are designed to maintain vouchers (including types) for long-term preservation, and increasingly they implement institutionalized workflows that link DNA sequences to specimens and specimen metadata (Prendini et al., 2002). We strongly encourage the deposition of voucher specimens from field surveys and experimental studies in museum collections, as has been urged by the Entomology Collections Network (ECN) and required in many PhD programs. No matter how quickly new molecular techniques are developed or refined, there is no substitute for a reliable database of voucher specimens when it comes to ensuring the repeatability of biological research (Funk et al., 2005; Lendemer et al., 2020).

Our results show that species richness of the parasitic wasps that attack *Drosophila* is severely underestimated, and only a fraction of them have been described. In DROP, 38% of the OTUs are provisional species, and more than 46% of the named OTUs have synonyms. Remarkably, *Leptopilina heterotoma*, one of the world’s most studied parasitoids, has more than 20 synonyms! As is generally the case, the rate of species description and revision of *Drosophila* parasitoids lags far behind that with which molecular sequence data are generated. Ensuring a consistent application of OTU recognition is therefore essential. With DROP, researchers may ensure consistency in their application of scientific names, and that those names are valid, making the daunting process of describing *Drosophila* parasitoids more accurate and efficient. In addition to the collection of physical museum resources, a central role taxonomists play in DROP and its curation is that of fostering better integration of taxonomy with experimental and biodiversity research. Our intention is to perpetuate DROP beyond this introductory publication. We hope that experts in all areas of *Drosophila*-parasitoid biology and related fields will join us in this effort.

### Conclusion

Taxonomic confusion presents many obstacles in experimental and biodiversity studies. One way of addressing this impediment is to provide a reliable DNA library with traceable vouchers (Astrin et al., 2013). Compared to BOLD and GenBank, DROP is a small database that provides some advantages over an immense genetic database. For example, it is easier for the research community to have direct communication amongst themselves, when there is a strong focus on a few specific taxa (Weigand et al., 2019). A good database has to maintain good quality of molecular data, but even more challenging is to maintain quality of identification from different sources (Fontes et al., 2021). In a big database, setting up a universal standard that satisfied all the taxa and researchers desires is particularly challenging. The curated nature of DROP will allow us to make strong rules to govern this data and assure users of its fidelity. While GenBank and BOLD each perform some amount of curation, it could be difficult to agree on curators for the whole range of animal and plant species catalogued there. We developed DROP as a resource and platform for gathering and sharing reliable genomic sequence data for *Drosophila* parasitoids. We hope it will serve as a model for researchers working with organisms which present similar difficulties. While compiling DROP, we found that the high number of provisional *versus* named OTUs suggests that the diversity of parasitic wasps attacking *Drosophila* is greatly underestimated. With this in mind, DROP represents the start of an important knowledge base that will strengthen future studies of natural host-parasitoid interactions, population dynamics, biocontrol, and the impact of climate change on biodiversity and ecosystem services.

## Supporting information

DROP_Supplemental_material

## Acknowledgements

The idea of DROP project was developed during the 2018 Entomological Society of America conference, during the symposium “*Drosophila* parasitoids: from molecular to ecosystem level”. We thank Dr. Elijah Talamas for valuable comments on earlier drafts and Dr. Vid Bakovic for genomic consultation on this project. We also thank Chris Jeffs for providing some Australian field samples. We are thankful for funding support from the Czech Science Foundation (17-27184Y). Additional fund for sequencing was provided by MLB, OTL, and PPAS. Mention of trade names or commercial products in this publication is solely for the purpose of providing specific information and does not imply recommendation or endorsement by the USDA. USDA is an equal opportunity provider and employer.

## Data Accessibility

The DROP database is freely accessible at Zenodo depository (http://doi.org/10.5281/zenodo.4519656). Sequences from GenBank and BOLD, all compiled in BOLD, DROP project, DS-DROPAR dataset <u>dx.doi.org/10.5883/DS-DROPAR.</u> New sequences have been deposited in BOLD, DROP project (datasets: DS-LABS <u>dx.doi.org/10.5883/DS-LABS</u> and DS-AUSPTOID <u>dx.doi.org/10.5883/DS-AUSPTOID</u>).

## Author Contributions

The initial project idea was originated by C-HL, MLB, JH, MM, TS, JV, SG, and PPAS. Molecular work was conducted by C-HL, SS, ML, AJ, and AD. BOLD and GenBank data was harvested by TAE and C-HL. Figures were made by AL and C-HL. Laboratory and field sample preparations were conducted by MTK, YC, TS, MM, SG, JV, EG, MG, XW, KM, KMD, PA, NAP, MT, JJB, MP, FMJ, WDT, JSD, BW, OTL, PPAS, JL and AL. Taxonomic concepts and interpretations were conducted by RRK, MLB, CH-L, PG, and SEM. DROP database was built by JH and C-HL. All authors contributed to review and final revisions of the manuscript, which was written primarily by C-HL, MLB and JH.

