## Supplementary material for "DROP: Molecular voucher database for identification of *Drosophila* parasitoids": DROP_Supplemental_material: DROP_Supp_Documents_5.10.21.pdf

### Supplemental Materials

#### *Procedure for obtaining laboratory strain sequences*

DNA was extracted from three legs of each female by using 96-well plates which are preserved in a 2D barcode tissue tubes. The rest of the body of the individual was dry-mounted and assigned with another 2D USNM barcode that links to their DNA extraction. The fragment of the COI barcode was systematically amplified for each specimen by a robot. CO1: We amplified wasp COI DNA using Promega's GoTaq HotStart Master Mix. Starting with primers LCO1490 (GGTCAACAAATCATAAAGATATTGG Folmer) and Nancy (CCCGGTAAAATTAAAATATAAACTTC, AKA UBC9 or C1-N-2191R, Simon 1994), we ran 10 µl reactions, with the following reagents: 3.3 µl H20, 5 µl 2x GoTaq HotStart Master Mix, 0.3 µl each primer (at 10 µM concentration), and 0.1 µl BSA (at 20 mg/ml). 1 µl of undiluted template was added to each reaction. We used the following thermocycler profile: an initial denaturation of 95°C for 7 min, followed by 35 cycles of 95°C for 30s, 50°C for 30s, 72°C for 45s, with a final elongation step of 72°C for 1:30 min. Some wasps had a long poly A(T) region towards the 5' end of our region of interest that prohibited getting good overlapping forward and reverse reads. In these cases, we used 3 different forward primers that were all ~ 170-220 bp downstream of LCO1490 (and downstream of the poly A(T) region: RonMWaspdeg (GGWTCWCCWGATATAKCWTTTCC Smith et al 2009), UBC6 (GGAGGATTTGGAAATTGATTAGTGCC, AKA C1-J-1718F, Simon 1994), and Cl13 (ATAATTTTTTTATAGTTATACC, Clary and Wolstenholme 1985). We paired all these primers with Nancy, and we generally amplified each sample for each primer pair and sequenced the amplification that worked best. Cl13 gives the longest read (by 36 bp over UBC6 and 68 over Ron) followed by UBC6 (by 32 bp over Ron). For sequencing, we used 10 µl reactions that contained 6.5 µl H20, 1.75 µl Big Dye Buffer (BD Buffer), 0.25 µl primer (at 10 µM), 0.5 µl Big Dye, and 1 µl of PCR product. We ran the following thermocycler profile: an initial denaturation of 96°C for 1 min, followed by 30 cycles of 96°C for 30s, 50°C for 30s, and 60°C for 1:30 min. Beyond standard COI barcoding, additional genes were sequenced for molecular systematic purposes in come specimens, including ITS2. The primers ITS2F (5'-ATTCCCGGACCACGCCTGGCTGA-3') and ITS2R (5'-TCCTCCGCTTATTGATATGC-3') were used to amplify ITS2 region (White et al. 1990). Thermal cycling conditions included an initial denaturation step at 94 °C for 4 min, followed by 35 cycles of 94 °C for 30 s, 47 °C for 30 s and 72 °C for 40 s, with a final step of 10 min at 94 °C.

Aligned COI sequences were used to construct a Neighbor Joining tree using ClustalW v.2.1 (Larkin et al 2007), implemented in the EMBL-EBI web interface (Madeira et al 2019), using default tree-building parameters (distance correction, gap exclusion, and PIM set as -false). The Newick file was imported into FigTree v.1.4.4 (https://github.com/rambaut/figtree/) and manually rooted on the outgroup. The phylogeny was further annotated in inkscape (https://inkscape.org/).

***Step by step instruction for install SQLiteStudio and opening DROP***

1. To install SQLite3 check the latest release version and download (https://www.sqlite.org/) for your specific operating system, e.g., Windows, Linux, or Mac.
2. The download file is in ZIP format. Unzip and run SQLite
3. After installing SQLite3, Go to SQLiteStudio (https://github.com/pawelsalawa/sqlitestudio/releases) check your operating system and download. (SQLite3 will actually be run through SQLiteStudio).
4. Go to (http://doi.org/10.5281/zenodo.4519656) and download DROPsqlite3 file (this file contains all of the sequences and metadata in the database in a single file)
5. Run the SQLiteStudio program.
6. Go to file-> open-> indicate directory where DROP database is saved.
7. You can now visualize, use and explore the DROP database for your intended application. You can explore and modify the existing queries.

**Supplemental Tables**

Table S1: List of parasitoid species with potential BIN-ID conflicts, either associated with more than one BINs or with multiple species sharing the same BIN. Green-color cells (AAJ9287) associated with both *Asobara japonica* and *A. tabida*. Orange-color cells (ACD4029, ACD4086, ACD2049, ACD4028) associated with both *Ganaspis brasiliensis* and *G. xanthopoda*.

| **Genus** | **Species** | **BIN** |
|---|---|---|
| *Asobara* | *Asobara japonica* | BOLD:ACB7213 |
| | | BOLD:AAJ9287 |
| | | BOLD:ACD3963 |
| *Asobara* | *Asobara mesocauda* | BOLD:ADX7763 |
| | | BOLD:ADK0178 |
| | | BOLD:ADU9416 |
| *Asobara* | *Asobara pleuralis* | BOLD:ACD4008 |
| | | BOLD:ACB8130 |
| *Asobara* | *Asobara rufescens* | BOLD:AAU8583 |
| | | BOLD:ACQ2439 |
| *Asobara* | *Asobara tabida* | BOLD:ACD4234 |
| | | BOLD:AAJ9287 |
| *Aspilota* | *Aspilota angusta* | BOLD:AAV4382 |
| | | BOLD:ABA5938 |
| *Ganaspis* | *Ganaspis brasiliensis* | BOLD:ACD4029 |
| | | BOLD:ACD4086 |
| | | BOLD:ABX2049 |
| | | BOLD:ACD4028 |
| *Ganaspis* | *Ganaspis xanthopoda* | BOLD:ACR2271 |
| | | BOLD:ACD4028 |
| | | BOLD:ABX2049 |
| | | BOLD:ACD4029 |
| | | BOLD:ADJ7530 |
| | | BOLD:ACD4086 |
| *Leptopilina* | *Leptopilina victoriae* | BOLD:ACB8261 |
| | | BOLD:ACB8262 |

Table S2: List of institutions and their abbreviations included in DROP.

| **Institute Full Name** | **Abbreviation** |
|---|---|
| National Museum of Natural History, Smithsonian Institution, Washington DC | USNM |
| Natural History Museum, London | BMNH |
| Swedish University of Agricultural Sciences | SUAS |
| Systematic Entomology Hokkaido University | SEHU |
| The Triplehorn Insect Collection, The Ohio State University | OSUC |
| Zoology, Chinese Academy of Sciences | IZCAS |
| Natural Biodiversity Center, Leiden, Netherlands | RMNH |
| Australian National Insect Collection, CSIRO, Canberra | ANIC |
| Zoological Museum, University of Copenhagen | ZMUC |
| New Zealand Arthropod Collection | NZAC |
| Zoologiska Museet, Lund University, Lund | MZLU |
| Museum National d'Histoire naturelle, Paris | MNHN |
| Zoological Institute RAS of St Petersburg, Russia | ZISP |
| Center of Biodiversity Genomics | CBG |
| Bernice P. Bishop Museum Honolulu | BPBM |
| Entomology Research Collection, UC Riverside | UCR |
| Institute of Insect Sciences, Zhejiang University, Hangzhou, China | ZHU |
| Naturhistorisches Museum Wien, Austria | NMWA |
| Zoologisches Muesum, Museum fur Naturkunde an der Humboldt-Universitat zu Berlin, G.D.R. | MNB |
| Zoologisches Sammlung des Bayerischen Staates, Munich | ZSBM |
| Bishop Museum | BISHOP |
| The Czech Academy of Sciences | CAS |
| Institute for Sustainable Plant Protection, National Research Council of Italy | CNR-IPSP |
| Ehime University | |
| Fujian Agriculture and Forest University | FAFU |
| Queensland Museum | QM |
| Texas A&M University | TAMU |
| The University of Arizona | |
| City University of New York | CUNY |
| Université Claude Bernard Lyon 1 | UCBL |
| Indian University of Bloomington | |
| University of Cambridge | |
| Université Côte d'Azur / Institut Sophia Agrobiotech | UCA / ISA |
| University Paris-Aclay | |
| United States department of Agriculture- Agricultural Research Services | USDA-ARS |
| Agriculture and Agri-Food Canada | AAFC |

Table S3: Laboratory strains listed in DROP.

| Strain_id | Strain_name | Laboratory | Alive | Species_id | DROP_Species_name |
|---|---|---|---|---|---|
| 1 | AcIC | Todd Schlenke | YES | 163 | *drop_ Aly_sp48* |
| 2 | AjJap | Todd Schlenke | YES | 165 | *drop_ Aly_sp50* |
| 3 | AtFr | Todd Schlenke | YES | 162 | *drop_ Aly_sp47* |
| 4 | TriCal | Todd Schlenke | YES | 159 | *drop_ Tri_sp44* |
| 5 | GxUg | Todd Schlenke | YES | 166 | *drop_ Gan_sp51* |
| 6 | GxHondu | Todd Schlenke | YES | 166 | *drop_ Gan_sp51* |
| 7 | GhFl | Todd Schlenke | YES | 168 | *drop_ Gan_sp53* |
| 8 | GhHaw | Todd Schlenke | YES | 167 | *drop_ Gan_sp52* |
| 9 | GbAtl | Todd Schlenke | YES | 167 | *drop_ Gan_sp52* |
| 10 | G2 | Todd Schlenke | YES | 167 | *drop_ Gan_sp52* |
| 11 | Lb17 | Todd Schlenke | YES | 173 | *drop_ Lep_sp58* |
| 12 | LcNet | Todd Schlenke | YES | 169 | *drop_ Lep_sp54* |
| 13 | LmAtl | Todd Schlenke | YES | 174 | *drop_ Lep_sp59* |
| 14 | Lh14 | Todd Schlenke | YES | 6 | *Leptopilina heterotoma* |
| 15 | LhSwe | Todd Schlenke | YES | 6 | *Leptopilina heterotoma* |
| 16 | LvHaw | Todd Schlenke | YES | 170 | *drop_ Lep_sp55* |
| 17 | LvPhil | Todd Schlenke | YES | 170 | *drop_ Lep_sp55* |
| 18 | LvUnk | Todd Schlenke | YES | 170 | *drop_ Lep_sp55* |
| 19 | LpIndo | Todd Schlenke | YES | 172 | *drop_ Lep_sp57* |
| 20 | LlJap | Todd Schlenke | YES | 171 | *drop_ Lep_sp56* |
| 21 | PachyPort | Todd Schlenke | YES | 160 | *drop_ PachyPort_sp45* |
| 22 | SFNS | Julien Varaldi | YES | 173 | *drop_ Lep_sp58* |
| 23 | Lb84 NS | Julien Varaldi | YES | 173 | *drop_ Lep_sp58* |
| 24 | Lb84 S | Julien Varaldi | YES | 173 | *drop_ Lep_sp58* |
| 25 | G495 NS | Julien Varaldi | YES | 173 | *drop_ Lep_sp58* |
| 26 | G495 S | Julien Varaldi | YES | 173 | *drop_ Lep_sp58* |
| 27 | NS1c | Julien Varaldi | YES | 173 | *drop_ Lep_sp58* |
| 28 | NS1bS | Julien Varaldi | YES | 173 | *drop_ Lep_sp58* |
| 29 | LhGot | Julien Varaldi | NO | 6 | *Leptopilina heterotoma* |
| 30 | GbHaw | Julien Varaldi | YES | 166 | *drop_ Gan_sp51* |
| 31 | GbVa | Julien Varaldi | YES | 166 | *drop_ Gan_sp51* |
| 32 | Td | Julien Varaldi | NO | | |
| 33 | Pv | Julien Varaldi | NO | | |
| 34 | Lg500 | Mariana Mateos | YES | 158 | *drop_Lg500_sp43* |
| 35 | GxHawM | Mariana Mateos | YES | 166 | *drop_ Gan_sp51* |

| | | | | | |
|---|---|---|---|---|---|
| 36 | G1FL | Mariana Mateos | YES | 168 | *drop_ Gan_sp53* |
| 37 | Lh14M | Mariana Mateos | YES | 6 | *Leptopilina heterotoma* |
| 38 | GxUgM | Mariana Mateos | NO | | |
| 39 | Lc03 | Mariana Mateos | YES | 175 | *drop_ Lep_sp60* |
| 40 | W2 | Mariana Mateos | YES | 177 | *drop_ Lep_sp62* |
| 41 | Lb17M | Mariana Mateos | NO | | |
| 42 | PV20 | Jan Hrcek | YES | 116 | *drop_Aso_sp8* |
| 43 | KHB | Jan Hrcek | YES | 116 | *drop_Aso_sp8* |
| 44 | 124PL | Jan Hrcek | NO | | |
| 45 | 179C | Jan Hrcek | YES | 116 | *drop_Aso_sp8* |
| 46 | 69B | Jan Hrcek | YES | 109 | *drop_Gan1_sp1* |
| 47 | 75B | Jan Hrcek | YES | 109 | *drop_Gan1_sp1* |
| 48 | 84BC | Jan Hrcek | YES | 109 | *drop_Gan1_sp1* |
| 49 | 96BKH | Jan Hrcek | YES | 109 | *drop_Gan1_sp1* |
| 50 | 106BPL | Jan Hrcek | YES | 110 | *drop_Fig124_sp2* |
| 51 | PHC1 | Jan Hrcek | NO | | |
| 52 | PHC2 | Jan Hrcek | NO | | |
| 53 | 111F | Jan Hrcek | YES | 110 | *drop_Fig124_sp2* |
| 54 | 161A | Jan Hrcek | YES | 110 | *drop_Fig124_sp2* |
| 55 | 126PH | Jan Hrcek | YES | 109 | *drop_Gan1_sp1* |
| 56 | 70AD | Jan Hrcek | YES | 117 | *drop_Dia127_sp9* |
| 57 | 66LD | Jan Hrcek | YES | 117 | *drop_Dia127_sp9* |
| 58 | 175A | Jan Hrcek | NO | | |
| 59 | Lh14G | Shubha Govind | YES | 6 | *Leptopilina heterotoma* |
| 60 | Lh14Space_Space | Shubha Govind | YES | 6 | *Leptopilina heterotoma* |
| 61 | Lh14Ground_Ground | Shubha Govind | YES | 6 | *Leptopilina heterotoma* |
| 62 | LvNet | Shubha Govind | YES | 170 | *drop_ Lep_sp55* |
| 63 | Lh14 aurum18 | Shubha Govind | YES | 6 | *Leptopilina heterotoma* |
| 64 | Tk (Lh_Kimura) | Shubha Govind | YES | 6 | *Leptopilina heterotoma* |
| 65 | RhoTs | Shubha Govind | NO | | |
| 66 | G486_Govind | Shubha Govind | YES | 173 | *drop_ Lep_sp58* |
| 67 | TdG | Shubha Govind | YES | 159 | *drop_ Tri_sp44* |
| 68 | KS (Gb Kimura) | Shubha Govind | YES | 166 | *drop_ Gan_sp51* |
| 69 | LbG486G | Shubha Govind | YES | 174 | *drop_ Lep_sp59* |

| | | | | | |
|---|---|---|---|---|---|
| 70 | JCU | Jan Hrcek | NO | | |
| 71 | STL6 | Dan Tracey | YES | 115 | *drop_STL_sp7* |
| 72 | Guad1 | Dan Tracey | YES | 170 | *drop_ Lep_sp55* |
| 73 | GREN6 | Dan Tracey | YES | 170 | *drop_ Lep_sp55* |
| 74 | STK1 | Dan Tracey | YES | 115 | *drop_ STL_sp7* |
| 75 | Lc01 | Mariana Mateos | YES | 176 | *drop_ Lep_sp61* |
| 76 | PachyPort_old | Todd Schlenke | YES | 160 | *drop_ PachyPort_sp45* |
| 77 | TriCal_old | Todd Schlenke | YES | 159 | *drop_ Tri_sp44* |
| 78 | L.japfor_Kimura | Masahito Kimura | NO | | |
| 79 | L.japjap_Kimura | Masahito Kimura | NO | | |
| 80 | L.longipes_Kimura | Masahito Kimura | NO | | |
| 81 | L.pacifica_Kimura | Masahito Kimura | NO | | |
| 82 | L.ryukyuensis_Kimura | Masahito Kimura | NO | | |
| 83 | L.tsushimaensis_Kimura | Masahito Kimura | NO | | |
| 84 | L.victoriae_Kimura | Masahito Kimura | NO | | |
| 85 | Carton 302-1 | Yves Carton | NO | | |
| 87 | G.brasiliensis_Kimura | Masahito Kimura | NO | | |
| 88 | G.brasiliensis_Danne | USDA-ARS quarantine | YES | 19 | *Ganaspis brasiliensis* |
| 89 | L. bNEA | Kent Daane | NO | | |
| 90 | L.bORI | Kent Daane | NO | | |
| 91 | L.hNEA | Kent Daane | NO | | |
| 92 | L.jORI | USDA-ARS quarantine | YES | 13 | *Leptopilina japonica* |
| 93 | L.vA_old | Shubha Govind | YES | 170 | *drop_Lep_sp55* |
| 94 | L.vD_old | Shubha Govind | YES | 170 | *drop_Lep_sp55* |
| 95 | variant_old | Shubha Govind | NO | | |
| 96 | LbKEN_old | Todd Schlenke | NO | | |
| 97 | LbFR_old | Todd Schlenke | NO | | |
| 98 | Lb17_old | Todd Schlenke | NO | | |
| 99 | LbG486_old | Todd Schlenke | NO | | |
| 102 | SevernMD | Jeff Leips | NO | | |

| | | | | | |
|---|---|---|---|---|---|
| 103 | L.japjap_Kimura_2 | Masahito Kimura | NO | | |
| 104 | L.heterotoma_Kimura | Masahito Kimura | NO | | |
| 105 | L.ryukyuensis_Kimura_2 | Masahito Kimura | NO | | |
| 106 | A.japonica_Kimura_1 | Masahito Kimura | NO | | |
| 107 | A. japonica_Kimura_2 | Masahito Kimura | NO | | |
| 108 | A. japonica_Kimura_3 | Masahito Kimura | NO | | |
| 109 | A. rufescens_Kimura | Masahito Kimura | NO | | |
| 110 | A. rossica_Kimura | Masahito Kimura | NO | | |
| 111 | A. pleuralis_Kimura | Masahito Kimura | NO | | |
| 112 | L. b 173-1 | Yves Carton | NO | | |
| 113 | L. b G 301-1 | Marylene Poirie | YES | 173 | *drop_ Lep_sp58* |
| 114 | L. b L104 (G483) | Yves Carton | NO | | |
| 115 | L. b France I | Yves Carton | NO | | |
| 116 | L. b Nasrallah_1978 | Yves Carton | NO | | |
| 117 | L. b Olbia | Yves Carton | NO | | |
| 118 | L. b Pont de la May | Yves Carton | NO | | |
| 119 | L. b Nasrallah (G317) | Yves Carton | NO | | |
| 120 | L. b L104 (G346) | Yves Carton | NO | | |
| 121 | L. b Nasrallah_1982 | Yves Carton | NO | | |
| 122 | L. b Nasrallah_1985 | Yves Carton | NO | | |
| 123 | L. b G483 (G346) | Yves Carton | NO | | |
| 124 | G. xanthopoda G 302-1 | Yves Carton | NO | | |
| 125 | L. b G486_Carton | Marylene Poirie | YES | 173 | *drop_ Lep_sp58* |
| 126 | L. b G486 (ISy) | Marylene Poirie | YES | 173 | *drop_ Lep_sp58* |
| 127 | L. b G 431 (ISm) | Marylene Poirie | YES | 173 | *drop_ Lep_sp58* |
| 128 | L. b G 397 | Yves Carton | NO | | |
| 129 | A. tabida 490 | Yves Carton | NO | | |

| | | | | | |
|---|---|---|---|---|---|
| 130 | L. b G 464 | Yves Carton | NO | | |
| 131 | L. b G 495 | Yves Carton | NO | | |
| 132 | L. freyae G499 | Yves Carton | NO | | |
| 133 | L. freyae G519 | Yves Carton | NO | | |
| 134 | L. orientalis G552-1 | Yves Carton | NO | | |
| 135 | L. guineaensis G510 | Yves Carton | NO | | |
| 136 | L. victoriae | Yves Carton | NO | | |
| 137 | L. b Lyon | Yves Carton | NO | | |
| 138 | L. h Lyon | Yves Carton | NO | | |
| 139 | L. h Saint Foy | Yves Carton | NO | | |
| 140 | L. victoriae G513.2 | Yves Carton | NO | | |
| 141 | L. victoriae G504.1 | Yves Carton | NO | | |
| 142 | L. guineaensis G500 | Yves Carton | NO | | |
| 143 | L. guineaensis G501 | Yves Carton | NO | | |
| 144 | L. orientalis G504.2 | Yves Carton | NO | | |
| 145 | NSRef | Frank Jiggins | YES | 173 | *drop_ Lep_sp58* |
| 146 | G486_Jiggins | Frank Jiggins | YES | 173 | *drop_ Lep_sp58* |
| 147 | Lh548 | Marylène Poirié | YES | 6 | *Leptopilina heterotoma* |
| 148 | Aph1Atl | Todd Schlenke | NO | | |
| 149 | Pac1Atl | Todd Schlenke | NO | | |
| 150 | Tri1Fr | Todd Schlenke | NO | | |
| 151 | Lb17G | Shubha Govind | YES | 173 | *drop_ Lep_sp58* |
| 152 | LhNY | Shubha Govind | YES | 6 | *Leptopilina heterotoma* |
| 153 | Gx | Shubha Govind | NO | | |
| 154 | GbVA | Shubha Govind | YES | 166 | *drop_ Gan_sp51* |
| 155 | GbHAW | Shubha Govind | YES | 166 | *drop_ Gan_sp51* |
| 156 | SAG_Nopal | Mariana Mateos | NO | | |
| 157 | Lb17_Lindsey | Amelia Lindsey | YES | 173 | *drop_ Lep_sp58* |

Table S4: *Drosophila* parasitoid whole-genome sequences included in DROP. For additional details, see DROP.

| Genus | Species_Name | Species_id | Genome_id | Voucher_id | GenBank_id |
|---|---|---|---|---|---|
| *Ganaspis* | *drop_Gan_sp51* | 166 | 8 | 868 | GCA_009823575.1 |
| *Ganaspis* | *brasiliensis* | 19 | 16 | 872 | SRX8882993 |
| *Ganaspis* | *brasiliensis* | 19 | 17 | 871 | SRX8882992 |
| *Ganaspis* | *drop_ Gsp1_sp67* | 182 | 15 | 873 | SRX8882994 |
| *Ganaspis* | *drop_ Gsp2_sp68* | 183 | 14 | 874 | SRX8882995 |
| *Ganaspis* | *drop_ Gsp50_sp66* | 181 | 9 | 869 | GCA_011057455.1 |
| *Leptolamina* | *ponapensis* | 48 | 13 | 875 | SRX8883008 |
| *Leptopilina* | *drop_Lep_sp58* | 173 | 5 | 865 | GCA_011634795.1 |
| *Leptopilina* | *drop_Lep_sp58* | 173 | 6 | 866 | GCA_003121605.1 |
| *Leptopilina* | *boulardi* | 4 | 12 | 876 | SRX8883009 |
| *Leptopilina* | *clavipes* | 5 | 7 | 867 | GCA_001855655.1 |
| *Leptopilina* | *heterotoma* | 6 | 1 | 861 | GCA_010016045.1 |
| *Leptopilina* | *heterotoma* | 6 | 2 | 862 | GCA_009602685.1 |
| *Leptopilina* | *heterotoma* | 6 | 3 | 863 | GCA_009026005.1 |
| *Leptopilina* | *heterotoma* | 6 | 4 | 864 | GCA_009025955.1 |
| *Leptopilina* | *japonica japonica* | 13 | 11 | 877 | SRX8883011 |

Table S5**:** *Drosophila* parasitoid transcriptome data included in DROP.

| Genus | Species_Name | Strain_id | Transcriptome_id | Voucher_id | Genbank_id |
|---|---|---|---|---|---|
| *Leptopilina* | *drop_Lep_sp58* | 126 | 2 | 858 | 2183568 |
| *Leptopilina* | *drop_Lep_sp58* | 127 | 3 | 859 | 2183567 |
| *Leptopilina* | *drop_Lep_sp58* | 151 | 8 | 882 | 15642271 |
| *Leptopilina* | *drop_Lep_sp58* | 151 | 9 | 883 | 15642270 |
| *Leptopilina* | *heterotoma* | 147 | 1 | 857 | 2183569 |
| *Leptopilina* | *heterotoma* | 152 | 5 | 884 | 2046288 |
| *Leptopilina* | *heterotoma* | 61 | 6 | 880 | 11581553 |
| *Leptopilina* | *heterotoma* | 60 | 7 | 881 | 11662592 |
| *Leptopilina* | *drop_Lep_sp58* | 11 | 10 | 908 | GAJA00000000.1 |
| *Leptopilina* | *heterotoma* | 14 | 11 | 909 | GAJC00000000.1 |
| *Ganaspis* | *drop_Gan_sp53* | 7 | 12 | 910 | GAIW00000000.1 |
| *Leptopilina* | *heterotoma* | 147 | 13 | 915 | 2183569 |
| *Leptopilina* | *drop_Lep_sp58* | 126 | 14 | 916 | 2183568 |
| *Leptopilina* | *drop_Lep_sp58* | 127 | 15 | 917 | 2183567 |

Table S6: *Drosophila* parasitoid proteomes data included in DROP

| Genus | Species_Name | Strain_id | Proteomes_id | Voucher_id | Assession_id |
|---|---|---|---|---|---|
| *Leptopilina* | *heterotoma* | 152 | 1 | 885 | PRIDE: PXD005639 |
| *Leptopilina* | *heterotoma* | 61 | 2 | 886 | PRIDE: PXD005632 |
| *Leptopilina* | *drop_Lep_sp58* | 27 | 3 | 911 | Upon request to |
| *Leptopilina* | *drop_Lep_sp58* | 11 | 4 | 912 | PRIDE: PDX023836 |
| *Leptopilina* | *heterotoma* | 14 | 5 | 913 | PRIDE: PDX023824 |
| *Ganaspis* | *drop_Gan_sp53* | 7 | 6 | 914 | PRIDE: PDX023825 |

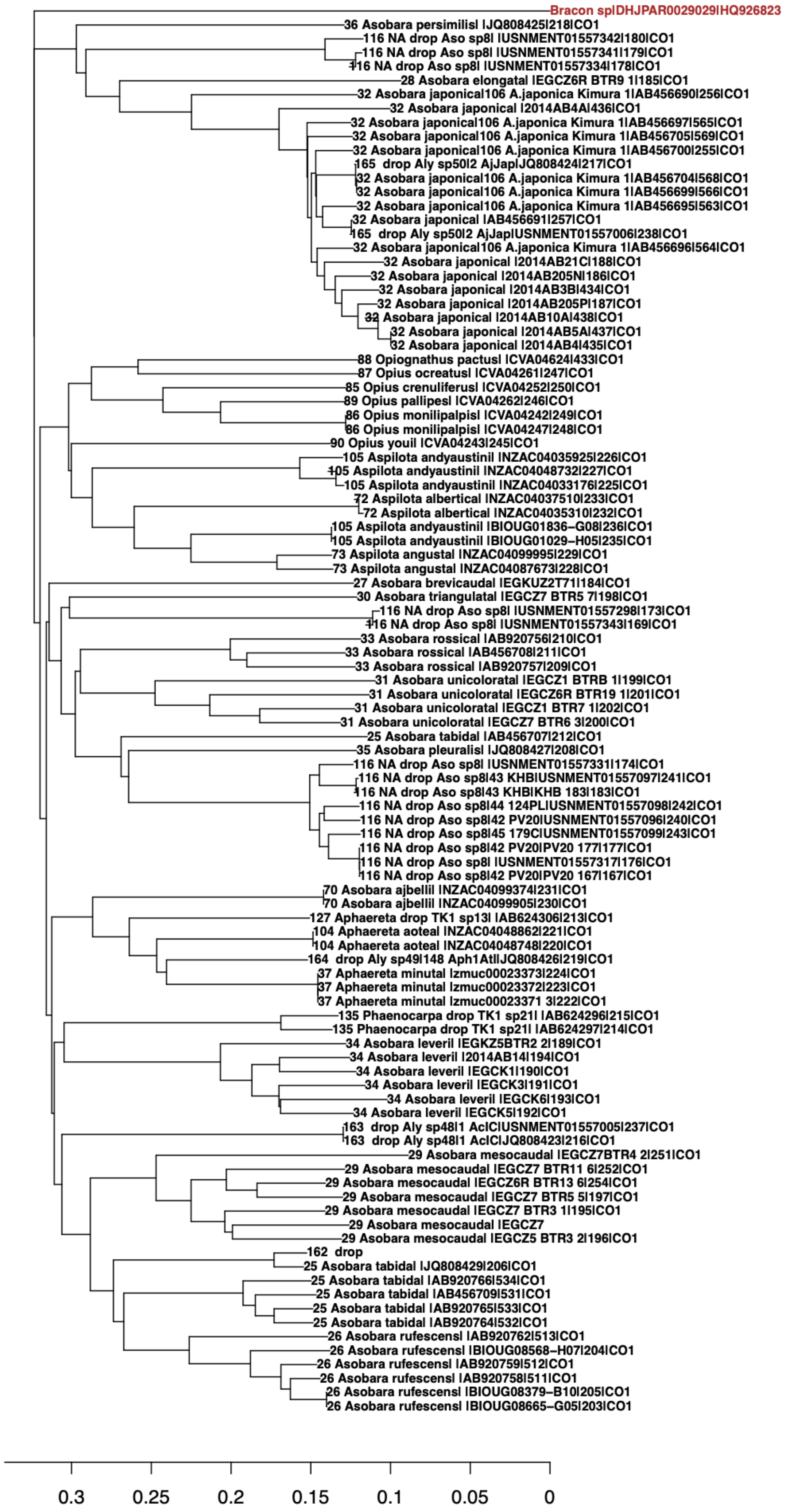
Bracon sp|DHJPAR0029029|HQ926823
36 Asobara persimilis| |JQ808425|218|CO1
116 NA drop Aso sp8| |USNMENT01557342|180|CO1
116 NA drop Aso sp8| |USNMENT01557341|179|CO1
116 NA drop Aso sp8| |USNMENT01557334|178|CO1
28 Asobara elongata| |EGCZ6R BTR9 1|185|CO1
32 Asobara japonica|106 A.japonica Kimura 1|AB456690|256|CO1
32 Asobara japonica| |2014AB4A|436|CO1
32 Asobara japonica|106 A.japonica Kimura 1|AB456697|565|CO1
32 Asobara japonica|106 A.japonica Kimura 1|AB456705|569|CO1
32 Asobara japonica|106 A.japonica Kimura 1|AB456700|255|CO1
165 drop Aly sp50|2 AjJap|JQ808424|217|CO1
32 Asobara japonica|106 A.japonica Kimura 1|AB456704|568|CO1
32 Asobara japonica|106 A.japonica Kimura 1|AB456699|566|CO1
32 Asobara japonica|106 A.japonica Kimura 1|AB456695|563|CO1
32 Asobara japonica| |AB456691|257|CO1
165 drop Aly sp50|2 AjJap|USNMENT01557006|238|CO1
32 Asobara japonica|106 A.japonica Kimura 1|AB456696|564|CO1
32 Asobara japonica| |2014AB21C|188|CO1
32 Asobara japonica| |2014AB205N|186|CO1
32 Asobara japonica| |2014AB3B|434|CO1
32 Asobara japonica| |2014AB205P|187|CO1
32 Asobara japonica| |2014AB10A|438|CO1
32 Asobara japonica| |2014AB5A|437|CO1
32 Asobara japonica| |2014AB4|435|CO1
88 Opiognathus pactus| |CVA04624|433|CO1
87 Opius ocreatus| |CVA04261|247|CO1
85 Opius crenuliferus| |CVA04252|250|CO1
89 Opius pallipes| |CVA04262|246|CO1
86 Opius monilipalpis| |CVA04242|249|CO1
86 Opius monilipalpis| |CVA04247|248|CO1
90 Opius youi| |CVA04243|245|CO1
105 Aspilota andyaustini| |NZAC04035925|226|CO1
105 Aspilota andyaustini| |NZAC04048732|227|CO1
105 Aspilota andyaustini| |NZAC04033176|225|CO1
72 Aspilota albertica| |NZAC04037510|233|CO1
72 Aspilota albertica| |NZAC04035310|232|CO1
105 Aspilota andyaustini| |BIOUG01836−G08|236|CO1
105 Aspilota andyaustini| |BIOUG01029−H05|235|CO1
73 Aspilota angusta| |NZAC04099995|229|CO1
73 Aspilota angusta| |NZAC04087673|228|CO1
27 Asobara brevicauda| |EGKUZ2T71|184|CO1
30 Asobara triangulata| |EGCZ7 BTR5 7|198|CO1
116 NA drop Aso sp8| |USNMENT01557298|173|CO1
116 NA drop Aso sp8| |USNMENT01557343|169|CO1
33 Asobara rossica| |AB920756|210|CO1
33 Asobara rossica| |AB456708|211|CO1
33 Asobara rossica| |AB920757|209|CO1
31 Asobara unicolorata| |EGCZ1 BTRB 1|199|CO1
31 Asobara unicolorata| |EGCZ6R BTR19 1|201|CO1
31 Asobara unicolorata| |EGCZ1 BTR7 1|202|CO1
31 Asobara unicolorata| |EGCZ7 BTR6 3|200|CO1
25 Asobara tabida| |AB456707|212|CO1
35 Asobara pleuralis| |JQ808427|208|CO1
116 NA drop Aso sp8| |USNMENT01557331|174|CO1
116 NA drop Aso sp8|43 KHB|USNMENT01557097|241|CO1
116 NA drop Aso sp8|43 KHB|KHB 183|183|CO1
116 NA drop Aso sp8|44 124PL|USNMENT01557098|242|CO1
116 NA drop Aso sp8|42 PV20|USNMENT01557096|240|CO1
116 NA drop Aso sp8|45 179C|USNMENT01557099|243|CO1
116 NA drop Aso sp8|42 PV20|PV20 177|177|CO1
116 NA drop Aso sp8| |USNMENT01557317|176|CO1
116 NA drop Aso sp8|42 PV20|PV20 167|167|CO1
70 Asobara ajbellii| |NZAC04099374|231|CO1
70 Asobara ajbellii| |NZAC04099905|230|CO1
127 Aphaereta drop TK1 sp13| |AB624306|213|CO1
104 Aphaereta aotea| |NZAC04048862|221|CO1
104 Aphaereta aotea| |NZAC04048748|220|CO1
164 drop Aly sp49|148 Aph1At|JQ808426|219|CO1
37 Aphaereta minuta| |zmuc00023373|224|CO1
37 Aphaereta minuta| |zmuc00023372|223|CO1
37 Aphaereta minuta| |zmuc00023371 3|222|CO1
135 Phaenocarpa drop TK1 sp21| |AB624296|215|CO1
135 Phaenocarpa drop TK1 sp21| |AB624297|214|CO1
34 Asobara leveri| |EGKZ5BTR2 2|189|CO1
34 Asobara leveri| |2014AB14|194|CO1
34 Asobara leveri| |EGCK1|190|CO1
34 Asobara leveri| |EGCK3|191|CO1
34 Asobara leveri| |EGCK6|193|CO1
34 Asobara leveri| |EGCK5|192|CO1
163 drop Aly sp48|1 AcIC|USNMENT01557005|237|CO1
163 drop Aly sp48|1 AcIC|JQ808423|216|CO1
29 Asobara mesocauda| |EGCZ7BTR4 2|251|CO1
29 Asobara mesocauda| |EGCZ7 BTR11 6|252|CO1
29 Asobara mesocauda| |EGCZ6R BTR13 6|254|CO1
29 Asobara mesocauda| |EGCZ7 BTR5 5|197|CO1
29 Asobara mesocauda| |EGCZ7 BTR3 1|195|CO1
29 Asobara mesocauda| |EGCZ7
29 Asobara mesocauda| |EGCZ5 BTR3 2|196|CO1
162 drop
25 Asobara tabida| |JQ808429|206|CO1
25 Asobara tabida| |AB920766|534|CO1
25 Asobara tabida| |AB456709|531|CO1
25 Asobara tabida| |AB920765|533|CO1
25 Asobara tabida| |AB920764|532|CO1
26 Asobara rufescens| |AB920762|513|CO1
26 Asobara rufescens| |BIOUG08568−H07|204|CO1
26 Asobara rufescens| |AB920759|512|CO1
26 Asobara rufescens| |AB920758|511|CO1
26 Asobara rufescens| |BIOUG08379−B10|205|CO1
26 Asobara rufescens| |BIOUG08665−G05|203|CO1
0.3
0.25
0.2
0.15
0.1
0.05
0

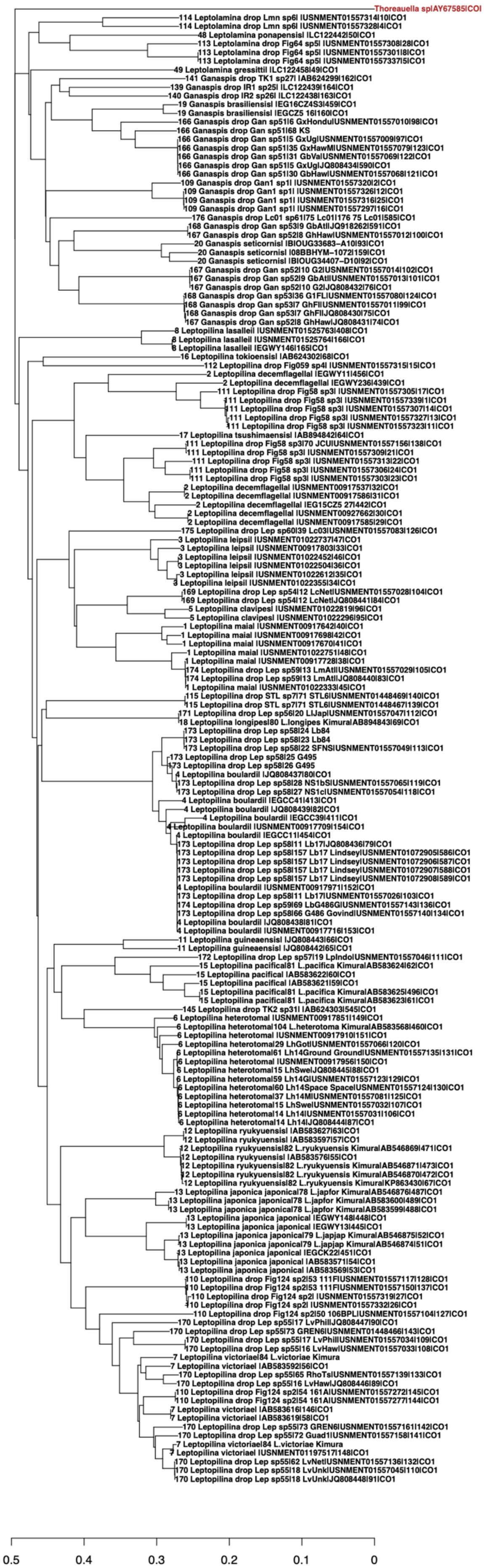

Thoreauella sp|AY67585|COI
114 Leptolamina drop Lmn sp6| |USNMENT01557314|10|CO1
114 Leptolamina drop Lmn sp6| |USNMENT01557328|4|CO1
48 Leptolamina ponapensis| |LC122442|50|CO1
113 Leptolamina drop Fig64 sp5| |USNMENT01557308|28|CO1
113 Leptolamina drop Fig64 sp5| |USNMENT01557301|8|CO1
113 Leptolamina drop Fig64 sp5| |USNMENT01557337|5|CO1
49 Leptolamina gressittii| |LC122458|49|CO1
141 Ganaspis drop TK1 sp27| |AB624299|162|CO1
139 Ganaspis drop IR1 sp25| |LC122439|164|CO1
140 Ganaspis drop IR2 sp26| |LC122438|163|CO1
19 Ganaspis brasiliensis| |EG16CZ4S3|459|CO1
19 Ganaspis brasiliensis| |EGCZ5 16|160|CO1
166 Ganaspis drop Gan sp51|6 GxHondu|USNMENT01557010|98|CO1
166 Ganaspis drop Gan sp51|68 KS
166 Ganaspis drop Gan sp51|5 GxUg|USNMENT01557009|97|CO1
166 Ganaspis drop Gan sp51|35 GxHawM|USNMENT01557079|123|CO1
166 Ganaspis drop Gan sp51|31 GbVa|USNMENT01557069|122|CO1
166 Ganaspis drop Gan sp51|5 GxUg|JQ808434|590|CO1
166 Ganaspis drop Gan sp51|30 GbHaw|USNMENT01557068|121|CO1
109 Ganaspis drop Gan1 sp1| |USNMENT01557320|2|CO1
109 Ganaspis drop Gan1 sp1| |USNMENT01557326|12|CO1
109 Ganaspis drop Gan1 sp1| |USNMENT01557316|25|CO1
109 Ganaspis drop Gan1 sp1| |USNMENT01557297|16|CO1
176 Ganaspis drop Lc01 sp61|75 Lc01|176 75 Lc01|585|CO1
168 Ganaspis drop Gan sp53|9 GbAt|JQ918262|591|CO1
167 Ganaspis drop Gan sp52|8 GhHaw|USNMENT01557012|100|CO1
20 Ganaspis seticornis| |BIOUG33683−A10|93|CO1
20 Ganaspis seticornis| |08BBHYM−1072|159|CO1
20 Ganaspis seticornis| |BIOUG34407−D10|92|CO1
167 Ganaspis drop Gan sp52|10 G2|USNMENT01557014|102|CO1
167 Ganaspis drop Gan sp52|9 GbAt|USNMENT01557013|101|CO1
167 Ganaspis drop Gan sp52|10 G2|JQ808432|76|CO1
168 Ganaspis drop Gan sp53|36 G1FL|USNMENT01557080|124|CO1
168 Ganaspis drop Gan sp53|7 GhFl|USNMENT01557011|99|CO1
168 Ganaspis drop Gan sp53|7 GhFl|JQ808430|75|CO1
167 Ganaspis drop Gan sp52|8 GhHaw|JQ808431|74|CO1
8 Leptopilina lasallei| |USNMENT01525763|408|CO1
8 Leptopilina lasallei| |USNMENT01525764|166|CO1
8 Leptopilina lasallei| |EGWY146|165|CO1
16 Leptopilina tokioensis| |AB624302|68|CO1
112 Leptopilina drop Fig059 sp4| |USNMENT01557315|15|CO1
2 Leptopilina decemflagella| |EGWY11|456|CO1
2 Leptopilina decemflagella| |EGWY236|439|CO1
111 Leptopilina drop Fig58 sp3| |USNMENT01557305|17|CO1
111 Leptopilina drop Fig58 sp3| |USNMENT01557339|1|CO1
111 Leptopilina drop Fig58 sp3| |USNMENT01557307|14|CO1
111 Leptopilina drop Fig58 sp3| |USNMENT01557327|13|CO1
111 Leptopilina drop Fig58 sp3| |USNMENT01557323|11|CO1
17 Leptopilina tsushimaensis| |AB894842|64|CO1
111 Leptopilina drop Fig58 sp3|70 JCU|USNMENT01557156|138|CO1
111 Leptopilina drop Fig58 sp3| |USNMENT01557309|21|CO1
111 Leptopilina drop Fig58 sp3| |USNMENT01557313|22|CO1
111 Leptopilina drop Fig58 sp3| |USNMENT01557306|24|CO1
111 Leptopilina drop Fig58 sp3| |USNMENT01557303|23|CO1
2 Leptopilina decemflagella| |USNMENT00917537|32|CO1
2 Leptopilina decemflagella| |USNMENT00917586|31|CO1
2 Leptopilina decemflagella| |EG15CZ5 27|442|CO1
2 Leptopilina decemflagella| |USNMENT00927662|30|CO1
2 Leptopilina decemflagella| |USNMENT00917585|29|CO1
175 Leptopilina drop Lep sp60|39 Lc03|USNMENT01557083|126|CO1
3 Leptopilina leipsi| |USNMENT01022737|47|CO1
3 Leptopilina leipsi| |USNMENT00917803|33|CO1
3 Leptopilina leipsi| |USNMENT01022452|46|CO1
3 Leptopilina leipsi| |USNMENT01022504|36|CO1
3 Leptopilina leipsi| |USNMENT01022612|35|CO1
3 Leptopilina leipsi| |USNMENT01022355|34|CO1
169 Leptopilina drop Lep sp54|12 LcNet|USNMENT01557028|104|CO1
169 Leptopilina drop Lep sp54|12 LcNet|JQ808441|84|CO1
5 Leptopilina clavipes| |USNMENT01022819|96|CO1
5 Leptopilina clavipes| |USNMENT01022296|95|CO1
1 Leptopilina maia| |USNMENT00917642|40|CO1
1 Leptopilina maia| |USNMENT00917698|42|CO1
1 Leptopilina maia| |USNMENT00917670|41|CO1
1 Leptopilina maia| |USNMENT01022751|48|CO1
1 Leptopilina maia| |USNMENT00917728|38|CO1
174 Leptopilina drop Lep sp59|13 LmAt|USNMENT01557029|105|CO1
174 Leptopilina drop Lep sp59|13 LmAt|JQ808440|83|CO1
1 Leptopilina maia| |USNMENT01022333|45|CO1
115 Leptopilina drop STL sp7|71 STL6|USNMENT01448469|140|CO1
115 Leptopilina drop STL sp7|71 STL6|USNMENT01448467|139|CO1
171 Leptopilina drop Lep sp56|20 LlJap|USNMENT01557047|112|CO1
18 Leptopilina longipes|80 L.longipes Kimura|AB894843|69|CO1
173 Leptopilina drop Lep sp58|24 Lb84
173 Leptopilina drop Lep sp58|23 Lb84
173 Leptopilina drop Lep sp58|22 SFNS|USNMENT01557049|113|CO1
173 Leptopilina drop Lep sp58|25 G495
173 Leptopilina drop Lep sp58|26 G495
4 Leptopilina boulardi| |JQ808437|80|CO1
173 Leptopilina drop Lep sp58|28 NS1bS|USNMENT01557065|119|CO1
173 Leptopilina drop Lep sp58|27 NS1c|USNMENT01557054|118|CO1
4 Leptopilina boulardi| |EGCC41|413|CO1
4 Leptopilina boulardi| |JQ808439|82|CO1
4 Leptopilina boulardi| |EGCC39|411|CO1
4 Leptopilina boulardi| |USNMENT00917709|154|CO1
4 Leptopilina boulardi| |EGCC11|454|CO1
173 Leptopilina drop Lep sp58|11 Lb17|JQ808436|79|CO1
173 Leptopilina drop Lep sp58|157 Lb17 Lindsey|USNMENT01072905|586|CO1
173 Leptopilina drop Lep sp58|157 Lb17 Lindsey|USNMENT01072906|587|CO1
173 Leptopilina drop Lep sp58|157 Lb17 Lindsey|USNMENT01072907|588|CO1
173 Leptopilina drop Lep sp58|157 Lb17 Lindsey|USNMENT01072908|589|CO1
4 Leptopilina boulardi| |USNMENT00917971|152|CO1
173 Leptopilina drop Lep sp58|11 Lb17|USNMENT01557026|103|CO1
174 Leptopilina drop Lep sp59|69 LbG486G|USNMENT01557143|136|CO1
173 Leptopilina drop Lep sp58|66 G486 Govind|USNMENT01557140|134|CO1
4 Leptopilina boulardi| |JQ808438|81|CO1
4 Leptopilina boulardi| |USNMENT00917716|153|CO1
11 Leptopilina guineaensis| |JQ808443|66|CO1
11 Leptopilina guineaensis| |JQ808442|65|CO1
172 Leptopilina drop Lep sp57|19 LpIndo|USNMENT01557046|111|CO1
15 Leptopilina pacifica|81 L.pacifica Kimura|AB583624|62|CO1
15 Leptopilina pacifica| |AB583622|60|CO1
15 Leptopilina pacifica| |AB583621|59|CO1
15 Leptopilina pacifica|81 L.pacifica Kimura|AB583625|496|CO1
15 Leptopilina pacifica|81 L.pacifica Kimura|AB583623|61|CO1
145 Leptopilina drop TK2 sp31| |AB624303|545|CO1
6 Leptopilina heterotoma| |USNMENT00917851|149|CO1
6 Leptopilina heterotoma|104 L.heterotoma Kimura|AB583568|460|CO1
6 Leptopilina heterotoma| |USNMENT00917910|151|CO1
6 Leptopilina heterotoma|29 LhGot|USNMENT01557066|120|CO1
6 Leptopilina heterotoma|61 Lh14Ground Ground|USNMENT01557135|131|CO1
6 Leptopilina heterotoma| |USNMENT00917956|150|CO1
6 Leptopilina heterotoma|15 LhSwe|JQ808445|88|CO1
6 Leptopilina heterotoma|59 Lh14G|USNMENT01557123|129|CO1
6 Leptopilina heterotoma|60 Lh14Space Space|USNMENT01557124|130|CO1
6 Leptopilina heterotoma|37 Lh14M|USNMENT01557081|125|CO1
6 Leptopilina heterotoma|15 LhSwe|USNMENT01557032|107|CO1
6 Leptopilina heterotoma|14 Lh14|USNMENT01557031|106|CO1
6 Leptopilina heterotoma|14 Lh14|JQ808444|87|CO1
12 Leptopilina ryukyuensis| |AB583627|63|CO1
12 Leptopilina ryukyuensis| |AB583597|57|CO1
12 Leptopilina ryukyuensis|82 L.ryukyuensis Kimura|AB546869|471|CO1
12 Leptopilina ryukyuensis| |AB583576|55|CO1
12 Leptopilina ryukyuensis|82 L.ryukyuensis Kimura|AB546871|473|CO1
12 Leptopilina ryukyuensis|82 L.ryukyuensis Kimura|AB546870|472|CO1
12 Leptopilina ryukyuensis|82 L.ryukyuensis Kimura|KP863430|67|CO1
13 Leptopilina japonica japonica|78 L.japfor Kimura|AB546876|487|CO1
13 Leptopilina japonica japonica|78 L.japfor Kimura|AB583600|489|CO1
13 Leptopilina japonica japonica|78 L.japfor Kimura|AB583599|488|CO1
13 Leptopilina japonica japonica| |EGWY148|448|CO1
13 Leptopilina japonica japonica| |EGWY13|445|CO1
13 Leptopilina japonica japonica|79 L.japjap Kimura|AB546875|52|CO1
13 Leptopilina japonica japonica|79 L.japjap Kimura|AB546874|51|CO1
13 Leptopilina japonica japonica| |EGCK22|451|CO1
13 Leptopilina japonica japonica| |AB583571|54|CO1
13 Leptopilina japonica japonica| |AB583569|53|CO1
110 Leptopilina drop Fig124 sp2|53 111F|USNMENT01557117|128|CO1
110 Leptopilina drop Fig124 sp2|53 111F|USNMENT01557150|137|CO1
110 Leptopilina drop Fig124 sp2| |USNMENT01557319|27|CO1
110 Leptopilina drop Fig124 sp2| |USNMENT01557332|26|CO1
110 Leptopilina drop Fig124 sp2|50 106BPL|USNMENT01557104|127|CO1
170 Leptopilina drop Lep sp55|17 LvPhil|JQ808447|90|CO1
170 Leptopilina drop Lep sp55|73 GREN6|USNMENT01448466|143|CO1
170 Leptopilina drop Lep sp55|17 LvPhil|USNMENT01557034|109|CO1
170 Leptopilina drop Lep sp55|16 LvHaw|USNMENT01557033|108|CO1
7 Leptopilina victoriae|84 L.victoriae Kimura
7 Leptopilina victoriae| |AB583592|56|CO1
170 Leptopilina drop Lep sp55|65 RhoTs|USNMENT01557139|133|CO1
170 Leptopilina drop Lep sp55|16 LvHaw|JQ808446|89|CO1
110 Leptopilina drop Fig124 sp2|54 161A|USNMENT01557272|145|CO1
110 Leptopilina drop Fig124 sp2|54 161A|USNMENT01557277|144|CO1
7 Leptopilina victoriae| |AB583616|146|CO1
7 Leptopilina victoriae| |AB583619|58|CO1
170 Leptopilina drop Lep sp55|73 GREN6|USNMENT01557161|142|CO1
170 Leptopilina drop Lep sp55|72 Guad1|USNMENT01557158|141|CO1
7 Leptopilina victoriae|84 L.victoriae Kimura
7 Leptopilina victoriae| |USNMENT01197517|148|CO1
170 Leptopilina drop Lep sp55|62 LvNet|USNMENT01557136|132|CO1
170 Leptopilina drop Lep sp55|18 LvUnk|USNMENT01557045|110|CO1
170 Leptopilina drop Lep sp55|18 LvUnk|JQ808448|91|CO1
0.5
0.4
0.3
0.2
0.1
0

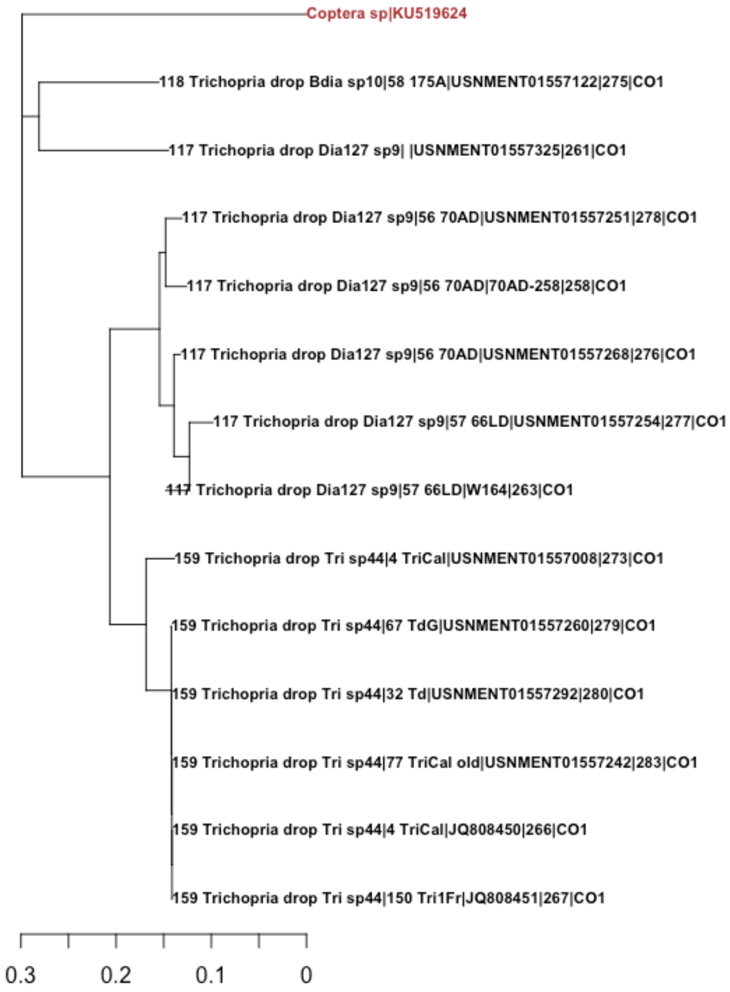


Figure S1: Neighbor-joining tree of Braconidae (a), Figitidae (b), and Diapriidae (c) indicating the potential species divergent for *Drosophila* lab strains based on CO1 sequences. The neighbor-joining trees were built only from the CO1 gene and therefore are not informative with regard to phylogeny. Outgroup is on the top (labeled in brown), the bottom axis indicates % genetic divergence. Node texts represent species or provisional species name of the lab strains, e.g.: 173 Leptopilina drop Lep sp58 (DROP species id, provisional species name) |27 NS1c (lab strain id, lab strain name) |USNMENT01557054 (Voucher number) |118 (DROP voucher id) |CO1 (genetic marker)

Figure S2: Structured Query Language (SQL) database structure of DROP. There are eight main linked tables: species, strain, voucher, sequence, genome, transcriptome, proteome, and reference (ref) tables connected in DROP. There are six additional tables linking the reference table with sequence, genome, transcriptome, proteome, species, and strain tables.

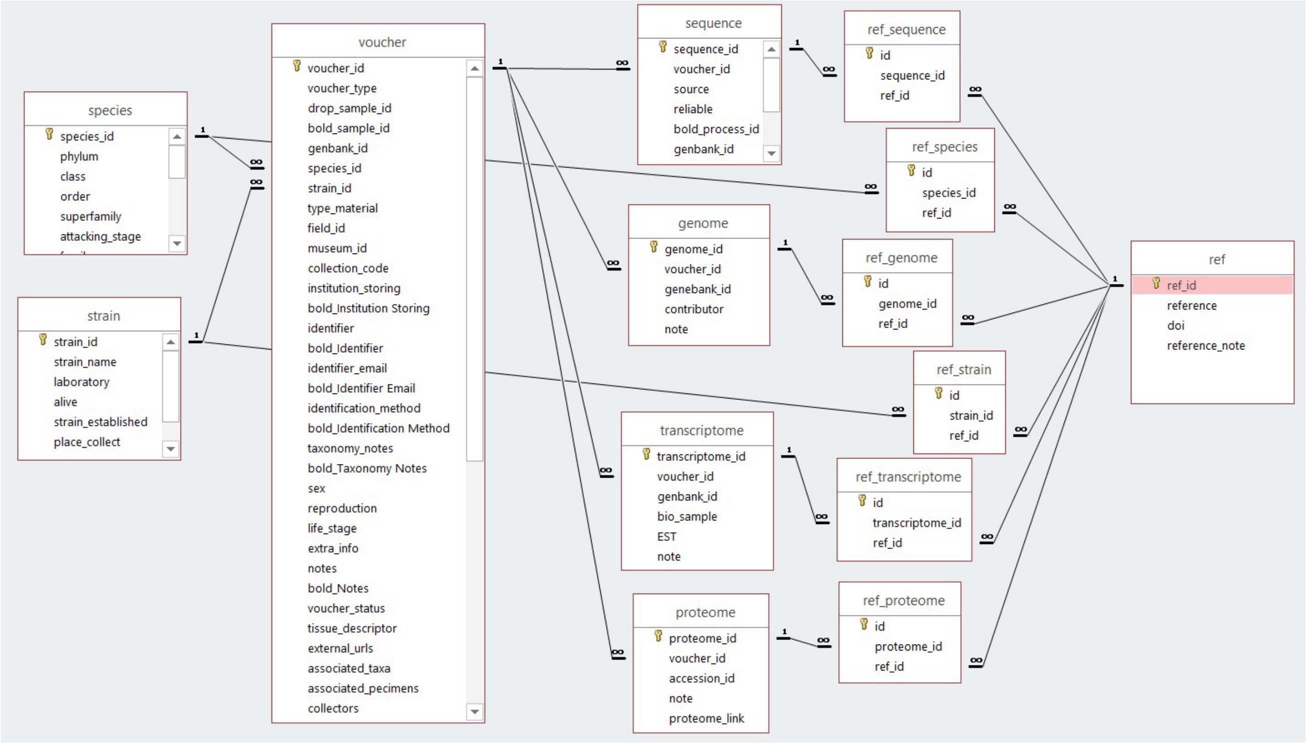
