## Supplementary material for "DROP: Molecular voucher database for identification of *Drosophila* parasitoids": DROP_Supplemental_material: README.md

# DROP
Molecular voucher database for identification of Drosophila parasitoids

Please cite as: ( https://doi.org/10.1101/2021.02.09.430471 ), later update to where it will be published.

Refer to the paper above for further details on the database. The database is manually curated, contact Chia-Hua Lue with updates and specimens to add.

Every release of the database contains the actual database file in sqlite3 format [ DROP.sqlite3 ] as well as a number of files generated from the database intended for easy use without delving into the primary database. These files are generated using views included in the main database file and can thus be easily modified to suit the needs of the users by adding fields and filtering the records. We use SQLiteStudio and sqlite3 programs (both free) to work with the database, but other programs can also work with the sqlite3 format.

Files generated from the database: 

DROP_sequences_fasta_COI_only.fas [ Reference COI sequences in fasta format. There are also separate files for individual superfamilies (Cynipoidea, Ichneumonoidea, Diaprioidea, and Chalcidoidea) ]

DROP_megatable.csv [ The core of the database (tables voucher, sequence, species and strain, but excluding genome, transcriptome, proteome and references tables) in a form of a very large table in a csv file. Intended for users not comfortable with the primary database format. From megatable view. ]

DROP_Species_catalog.csv  [ Currently a simple export of the species table. ]

DROP_Strains.csv [ List of strains with identifications. From strains view. ]

DROP_Genomes.csv [ List of genomes with identifications. From genomes view. ]

DROP_Transcriptomes.csv [ List of transcriptomes with identifications. From transcriptomes view. ]

DROP_Proteomes.csv [ List of transcriptomes with identifications. From proteomes view. ]

DROP_References.csv [ List of references - simple export of ref table ]

DROP_table_relationships.jpg   [ Picture of relationships between tables in the database ]

DROP_tables_field_description.xlsx [ Description of individual fields, with legend on fields match fields in BOLD (Barcoding of Life Database) ] 

-------------------------------------------------------------
How to generate fasta files from the database in Windows:

run sqlite3.exe [in same folder as database]
.open DROP.sqlite3
.output DROP_sequences_fasta_COI_only_Proctotrupoidea.fas

SELECT '>' || species.species_id || '_' || genus || '_' || species_name_only || '|' || COALESCE(strain.strain_id, '') || '_' || COALESCE(strain_name, '') || '|' || drop_sample_id || '|' || sequence_id || '|' || locus || x'0A' || sequence AS fasta
FROM sequence
LEFT OUTER JOIN voucher ON sequence.voucher_id = voucher.voucher_id
LEFT OUTER JOIN species ON voucher.species_id = species.species_id
LEFT OUTER JOIN strain ON voucher.strain_id = strain.strain_id
WHERE locus = 'CO1';

.quit

[you can copy the select query from the corresponding database view]
-------------------------------------------------------------

Shield: [![CC BY-SA 4.0][cc-by-sa-shield]][cc-by-sa]

This work is licensed under a
[Creative Commons Attribution-ShareAlike 4.0 International License][cc-by-sa].

[![CC BY-SA 4.0][cc-by-sa-image]][cc-by-sa]

[cc-by-sa]: http://creativecommons.org/licenses/by-sa/4.0/
[cc-by-sa-image]: https://licensebuttons.net/l/by-sa/4.0/88x31.png
[cc-by-sa-shield]: https://img.shields.io/badge/License-CC%20BY--SA%204.0-lightgrey.svg
