## Supplementary figures and images for "DROP: Molecular voucher database for identification of *Drosophila* parasitoids"

### DROP_table_relationships.jpg

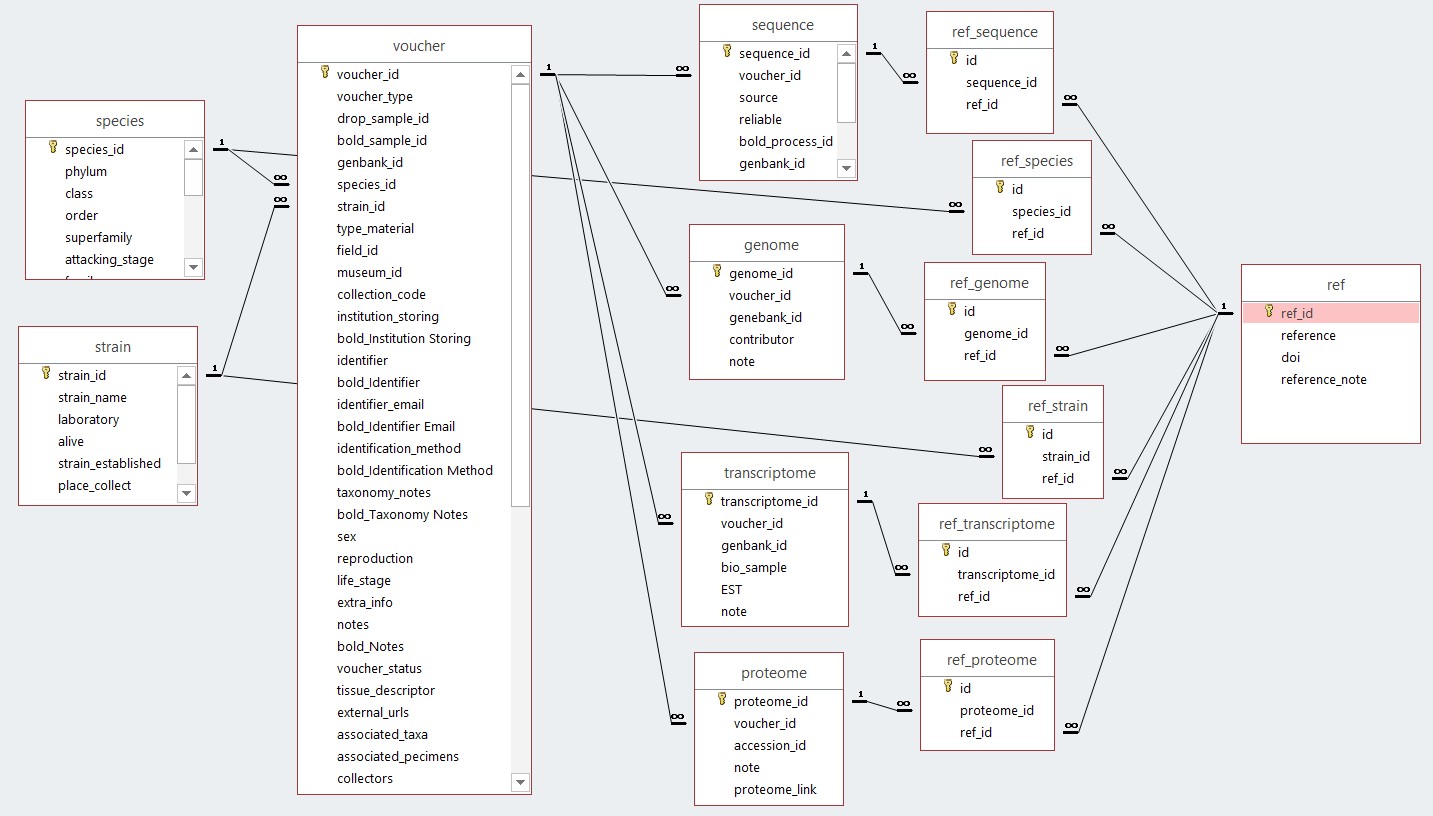
